# Robust Myocardial Regeneration After Selective Cardiomyocyte Loss Is Driven by Cardiac Stem Cell Activation Through the miR-221–p57 Axis

**DOI:** 10.64898/2026.07.05.736634

**Authors:** Eleonora Cianflone, Fabiola Marino, Mariangela Scalise, Andrew J Smith, Chiara Siracusa, Loredana Pagano, Claudia Quercia, Nadia Salerno, Assunta Di Costanzo, Giovanni Canino, Antonella De Angelis, Georgina M. Ellison-Hughes, Konrad Urbanek, Bernardo Nadal-Ginard, Daniele Torella

## Abstract

A central unresolved and highly contested question in cardiac biology is whether the adult mammalian heart, believed to have a very limited endogenous cardiomyocyte (CM) regenerative capacity, can be coaxed into an effective regenerative response after acute CM loss. Using TgMyh6^MCM^:R26^stop-DTA^ mice, we show that selective diffuse ablation of ∼15% of left ventricular CMs causes acute heart failure but is followed by complete structural and functional recovery within 28 days. Recovery is accomplished by robust generation of new mononucleated CMs, replacing ∼1/10 of the left ventricular CM compartment. This CM regeneration is produced by the activation of resident cardiac stem cells (CSCs), which exit quiescence, proliferate, produce new CMs, and subsequently return to quiescence. Depletion of the putative CSCs blocks repair, whereas transplantation of either clonogenic or primary CSCs through the systemic circulation fully restores myocardial regeneration and function, establishing that the CSCs home, nest and differentiate in the damaged myocardium and, therefore, are the main effectors of regeneration in this setting. Mechanistically, we show that miR-221-dependent repression of p57 governs the transition from quiescence--to activation—to differentiation—to quiescence of the CSCs, defining a reversible regulatory program which, under the proper conditions, endows the adult myocardium with robust CM regenerative competence.

## Introduction

The acute and/or chronic loss of cardiomyocytes (CMs), and the failure to efficiently replenish them, continues to be the number one cause of morbidity and mortality in developed societies even though the adult mammalian heart retains a measurable – but mostly underestimated--capacity for CM renewal throughout life^1–4^. The rate of this CM replenishment has been reported to be around 1% per year ^1, 2^, which highlights that this renewal is at best modest and ineffective in replacing most of the CM lost after a myocardial infarction. Furthermore, the source of this modest renewal has been highly contested. For practical purposes, biologically and clinically, the adult mammalian heart is still considered a post-mitotic organ without physiologically significant CM regenerative capability ^5–7^. This is in marked contrast to all other adult mammalian (including human) organs where parenchymal cell replenishment is mainly accomplished by an endogenous regenerative response entailing growth and differentiation of tissue-specific resident stem cells ^8–11^.

Although the heart is still viewed as a non-regenerative organ, resident cardiac stem/progenitor cells (CSCs) have been identified and characterized in the adult myocardium of all mammalian species studied as CD45^neg^/CD31^neg^ c-kit^pos^ and/or Sca-1^pos^ cells, which have been reproducibly proven by different groups to be clonogenic, self-renewing and multipotent capable of giving rise to CMs, vascular and connective cells *in vitro* and *in vivo* ^12–24^. Using a catecholamine injury model (single high-dose isoproterenol producing scattered microlesions predominantly in the sub-endocardium of the left ventricle’s (LV) apex), we provided evidence that the CSCs in the adult heart are both necessary and sufficient for functional cardiac regeneration and repair of acute cardiac damage: ablation of the CSCs abrogated recovery, whereas adoptive CSC transplantation into CSC-depleted hearts restored myocardial structure and function ^25, 26^. In contrast to the CSCs response to scattered microinjuries, segmental myocardial lesions, such as in myocardial infarction produced by a coronary artery occlusion, result in activation of the CSCs mainly in the lesion’s border with evident but limited regeneration at the boundaries of the damage, which fails to avoid and/or replace necrosis and scaring of the ischemic zone ^27–29^.

Over a decade ago, the data summarized above became controversial ^5, 7, 30–33^. First, the apparent inability of endogenous CSCs to effectively repair myocardial infarction became a source of scepticism in both the scientific and clinical communities ^5, 7, 34, 35^. Second, the retraction of several publications on CSCs by one of the early authors in the field, cast doubt on the broader body of independent research on these cells ^7, 34^. Third, a series of studies with important design limitations using site-specific recombination systems to lineage trace the cell fate of putative CSCs, concluded that in the adult myocardium, c-kit⁺ or Sca-1⁺ cardiac cells make a negligible, if any, contribution to the formation of new CMs ^30, 36–40^. These studies further argued that CMs are renewed almost exclusively by division of pre-existing CMs ^33, 36–40^. Not surprisingly, this plethora of negative findings further cemented the cell-static view of the heart and supported the idea that CSCs either do not exist or do not contribute meaningfully to CM renewal in adulthood, either under physiological conditions or after injury ^7, 33, 34^.

Consequently, the attention for regenerative approaches in the adult myocardium has focused on the characterization of a very large, and still growing, number of molecules and/or protocols which it is claimed can induce differentiated CMs re-entry to the cell cycle ^41–44^, to reprogram fibroblasts into CMs ^45, 46^, transplantation of autologous or allogenic *in vitro* generated CMs from pluripotent stem cells ^47–49^ or on the deployment of pluripotent stem cell-derived myocardial patches over the damaged region ^50–52^.

In the present study, we have addressed the capacity of the adult myocardium to renew its CMs by revisiting the role of CSCs using an approach established in classical bone marrow stem cell biology, implemented in a physiologically relevant injury context. We used transgenic mice to induce dose-dependent, CM-restricted Diphtheria Toxin A (DTA) activation, achieving a single-dose acute loss of approximately 15% of LV CMs which triggers an endogenous regenerative and functionally effective response. This response, in just one month, regenerates at least one tenth of the LV CMs. Using a combination of cell depletion and CSC transplantation, we show that endogenous CSCs are the main effectors of this CM renewal. Finally, we investigated the molecular mechanisms that govern injury-mediated CSCs activation cycle; that is, exit from quiescence, myocyte regeneration, and return to quiescence, identifying the miR-221–dependent repression of p57 as a key axis that orchestrates myocardial regeneration by the CSCs.

## Methods

### Animals

All animal handling and experimental procedures were performed according to the European Community guidelines (EC Council Directive 2010/63) and the Italian legislation on animal procedures (Decreto Legislativo D.Lgs 26/2014) and approved by Magna Graecia Institutional Review Boards on Animal Use and Welfare and by the Italian Ministry of Health, Section for veterinary ethics and animal care/use (Italian Ministry of Health, authorizations ex D.Lgs. January 27, 1992, n. 116; authorization number 368/2016-PR, released on 8 April 2016, extended on 25 November 2021; authorization number 823/2021PR released on: 27 October 2021). All efforts were made to minimize suffering, and the principles of Replacement, Reduction and Refinement (i.e., the “three Rs”) were applied to all experiments.

Mice were housed under controlled conditions at 25 °C, 50% relative humidity, and a 12 h light (6:00–18:00 h) and 12 h dark cycle, with water and food available ad libitum. All animals received human care, and all efforts were made to minimize animal suffering. Before any invasive procedure, the mice were anesthetized with intraperitoneal (i.p.) injections of tiletamine/zolazepam (80 mg/kg) or inhaled isoflurane (isoflurane, 1.5%; oxygen, 98.5%; Iso-Vet Piramal Healthcare).

Transgenic B6.FVB(129)-A1cfTg(Myh6-cre/Esr1*)1Jmk/J homozygous mice (hereafter TgMyh6^MCM^, Jackson lab Strain #:005657), expressing a tamoxifen (TAM)-inducible Cre Recombinase fusion protein (mER-Cre-mER) under the control of the cardiac α-myosin heavy chain (Myh6) promoter ^53^, were crossed with Cre-reporter homozygous mice carrying a mutation in the ROSA26 locus to express a floxed-STOP sequence in front of the Diphtheria Toxin A (DTA) gene (Strain R26^stop-DTA^, Jackson lab Strain #009669 or R26^eGFP-DTA^ Jackson lab Strain #032087, which display widespread expression of EGFP). TAM administration in double heterozygous TgMyh6^MCM^:R26^stop-DTA^ (hereafter TgMyh6-DTA) and TgMyh6^MCM^: R26^eGFP-DTA^ mice induces Cre-mediated excision of the STOP-cassette and activates DTA expression selectively in cardiomyocytes (CMs) leading to their ablation. Additionally, sex and age-matched C57BL/6J mice (Jackson Labs, stock number 000664) were used for experimental purposes. To assess CSC regenerative capacity *in vivo*, 4×10^5^ quiescent or activated CD45^neg^/CD31^neg^/c-kit^pos^ and/or Sca-1^pos^/GFP^pos^ CSCs, derived from *Tg-Myh6^MCM^:R26^eGFPstop-DTA^* donor mice two days after saline or TAM treatment, were injected though the tail vein into 5-FU–treated *Tg*Myh6-DTA mice 28 days post-TAM.

### Drug Administration

Twelve-sixteen weeks old double heterozygous male and female TgMyh6-DTA mice were administered with a single i.p. injection of TAM (Sigma-Aldrich) at dose of 0.5 mg, 0.25 mg or 0.125 mg. These TAM doses were established empirically through a trial-and-error experimental design that accounted for the unpredictable effects of mER-Cre-mER transgene copy number and expression among transgenic *Tg*Myh6^MCM^ mice. Two days after administration of 0.125mg TAM, TgMyh6-DTA mice were implanted with subcutaneous osmotic minipumps (ALZET) delivering either 5-Fluorouracil (5-FU, 15mg/Kg/day) or saline continuously for twenty-five days. At day twenty-eight, 5-FU-realizing mini-pumps were removed and 5-FU–treated TgMyh6-DTA mice were randomized to receive an intravenous injection of either saline or a GFP-labeled cardiac stem cell (CSC) clone. The animals were sacrificed 28 days after Saline/CSCs injections and the hearts were fixed in formalin or 4% paraformaldehyde (PFA) for immunohistochemistry analysis or dissociated to obtain a cardiac cell suspension.

### Myocyte necrosis analysis

To assess CM necrosis, twelve-sixteen weeks old double heterozygous TgMyh6-DTA mice, received an i.p. injection of 100 µg/100 µl of a monoclonal antibody against cardiac myosin (MF-20, ID: AB_2147781, DSHB) 2 h after 0.25 mg or 0.125 mg dose of TAM. All animals were sacrificed 48h after TAM injection, and heart were harvested and fixed in 4% PFA.

### Echocardiography

Echocardiography was performed as previously reported ^26, 54, 55^. Prior to echocardiography, mice were anesthetized with isoflurane. Unconscious mice were weighed and secured in a supine position on a temperature-controlled restraining board. Anesthesia was maintained with 1–2% isoflurane in oxygen delivered through a nose cone. Four-limb lead electrocardiograms (Vevo3100 and MP150, Biopac) were simultaneously recorded. All hair in the thoracic region was removed using a depilatory agent, and the area was cleaned with water. Ultrasound gel was applied to the thoracic region to improve sound wave transmission. All mice were maintained at heart rates > 400 bpm while images were recorded. Echocardiographic images were obtained with a Vevo 3100 system (Visualsonics, Inc.) equipped with a MX550D ultra-high frequency linear array transducer (22–55 MHz). The transducer was positioned in a stationary stand perpendicular to the mouse (in some cases, manual adaptations were needed for optimal imaging). In brief, a frame rate of >200 frames per minute was maintained for all B-mode and M-mode images. B-mode long-axis parasternal images were recorded when optimal views of the aorta, papillary muscle, and endocardium were visible. M-mode short-axis images were recorded at the level of the papillary muscles and the LV was bisected to obtain the optimal M-mode selection. Conventional echocardiographic measurements of the LV included ejection fraction (EF), fractional shortening (FS), end-diastolic dimension (EDD), end-systolic dimension (ESD), anterior and posterior wall thickness, and mass were obtained. For long-axis B-mode measurements, the endocardium was traced semi automatically beginning from the mitral valve and excluding the papillary muscle. EF and FS were calculated by software using standard computational methods. The apical 4-chamber view was acquired to assess mitral and tricuspid regurgitation. Advanced cardiac analysis (regional and global cardiac measurements) was assessed by speckle-tracking echocardiography (Vevo LAB analysis software; VisualSonics). Cardiac cycles were acquired digitally from the parasternal long-axis and mid-ventricular short-axis views for the assessment of radial, circumferential, and longitudinal systolic strain/velocity (in accordance with myocardial fiber orientation at varying levels of the LV wall) and time-to-peak systolic strain/velocity. Images selected for strain analysis had well-defined endocardium and epicardium borders and no substantial image artefacts. Image analysis was performed according to manufacturer’s instructions. The endocardium and epicardium were traced semi-automatically using VevoStrain software. The traces were manually adjusted to ensure adequate tracking of endocardium and epicardium borders. Velocity, displacement, strain, and strain rate were calculated for radial and longitudinal planes. In long-axis, the basal anterior-septum, mid-anterior-septum, apical anterior-septum, basal posterior wall, mid-posterior wall, and apical posterior segments were defined. In mid-ventricular short-axis, the anterior, anterior-septum, inferior-septum, inferior, posterior, and anterior-lateral segments were further delineated. Tissue contraction patterns were expressed as negative strain values for longitudinal and circumferential motion and positive values for radial strain. In each segment, peak systolic strain (%) and time-to-peak systolic strain (ms) were analyzed. Global average peak values for circumferential and longitudinal strain are reported. All measurements were performed offline using the VevoLab Analysis Software (FUJIFILM VisualSonics Inc)

### Mouse Cardiomyocyte nuclei isolation

Cardiomyocyte nuclei were isolated as previously described ^26, 56^. In brief, cardiac tissue was subjected to gentle mechanical disruption in a hypotonic buffer containing 0.31M sucrose, 10mM Tris-HCl, 5 mM CaCl2, 5 mM magnesium acetate, 2 mM EDTA, 0.5 mM EGTA and 1 mM DTT to selectively lyse the plasma membrane while preserving nuclear integrity. The lysate was then subjected to filtration and low-stringency centrifugation to remove large debris. Nuclei were subsequently enriched using a sucrose-based density separation procedure. Isolated nuclei were stained with anti-PCM1, anti-BrdU antibodies and analyzed by FACS.

### Mouse Cardiomyocyte isolation

Cardiomyocytes were isolated by enzymatic dissociation of hearts from twelve-sixteen weeks old TgMyh6-DTA and C57BL/6J mice. Hearts from TgMyh6-DTA and C57BL/6J mice were cannulated by aorta and hung on a retrograde perfusion system (Langendorff method), then perfused with enzyme-containing solutions as previously described ^26, 57^. A calcium-free solution was used to digest the hearts and prepared as follows: sodium chloride at a final concentration of 126 mM; glucose at a final concentration of 22 mM; HEPES at a final concentration of 24 mM; potassium chloride at a final concentration of 4.4 mM; magnesium dichloride at a final concentration of 5 mM; creatine at a final concentration of 5 mM; taurine at a final concentration of 20 mM; sodium pyruvate at a final concentration of 5 mM and Sodium dihydrogen phosphate at a final concentration of 1 mM. To increase the cardiomyocytes, yield 2,3-butanedione monoxime 10 mM was added to the solution. The solution was filtered through a 0.22 μm-pore filter into a sterile container and stored at 4 °C for up to 1 week. For heart isolation, a total amount of 50 ml of buffer was used. The cannulated hearts were first perfused with the calcium-free solution, followed by type II collagenase digestion in presence of Ca^2+^ 0.1 mM and then washed with a Ca^2+^ 0.1 mM solution. Hearts were then dissected into the left and right ventricles and the left and right atria. The atria were subsequently incubated on an orbital shaker in type II collagenase digestion solution supplemented with 0.1 mM Ca^2+^ for an additional 5 minutes at 37°C. Preparations were then filtered through a 100 μm cell strainer and viable cardiomyocytes were then allowed to sediment by gravity. The sedimentation procedure was repeated 3 times.

### Mouse Myocyte-depleted cardiac stem cell isolation

CSCs were isolated from the relative mouse hearts by enzymatic dissociation using a Langerdoff-modified apparatus or using gentle MACS Dissociator (Miltenyi Biotec). Briefly, for the Langerdoff-modified apparatus, the heart was excised, the aorta cannulated and hung on a retrograde perfusion system. Heart was perfused with collagenase type II dissolved HEPES-MEM (Sigma-Aldrich; Worthington) at 37 °C with 85% O2 and 15% N2, then the heart was removed from the apparatus, cut into small pieces, and the fragments shaken in re-suspension medium at 37 °C. CMs and myocyte-depleted small cardiac cells were separated by centrifugation. For gentle MACS-isolation, manufacturer instructions were followed to obtain myocyte-depleted cardiac small cells for FACS analysis. To obtain CSC-enriched CD45^neg^/CD31^neg^/c-kit^pos^ and/or Sca-1^pos^ cells, the MACS technology was used with direct CD45- and CD31-negative and then c-kit- and Sca-1-positive specific anti-mouse microbeads sorting (Miltenyi Biotec) ^23, 55^.

### Cell cultures

Isolated CSC-enriched CD45^neg^/CD31^neg^/c-kit^pos^ and/or Sca-1^pos^ cardiac cells were plated in gelatin-coated dishes in CSC growth medium containing Dulbecco’s modified Eagle’s medium-F12Ham’s (DMEM F-12, Gibco, Life Technologies) with insulin-transferrin-selenium (ITS 1%, Life Technologies), epidermal growth factor (EGF final medium concentration: 20 ng/ml, Peprotech), basal fibroblast growth factor (β-FGF final medium concentration: 10 ng/ml, Peprotech), and leukemia inhibitory factor (LIF, final medium concentration: 10 ng/ml, Miltenyi), and 1:1 ratio of Neurobasal medium (Gibco, Life Technologies) containing 37 mg of l-glutamine, B27 supplement (2%, Life Technologies) and N2 supplement (1%, Life Technologies), penicillin–streptomycin (1%, Life Technologies), Fungizone (0.1%, Life Technologies), and gentamicin (0.1%, Life Technologies) sterilized through a 0.22 µm pore filter into a sterile container, and store it at 4 °C for up to 3 weeks. The CSC growth medium was supplemented with 10% ESQ-FBS (Life Technologies). Cells were maintained at 37 °C in ambient O_2_ (21%) and 5% CO_2_. Media were replenished every 48 h and cells were passaged at a 1:4 ratio. The starvation medium was prepared as follow: Dulbecco’s modified Eagle’s medium-F12Ham’s implemented with 0.5% ESQ-FBS, penicillin–streptomycin (1%), Fungizone (0.1%), and gentamicin (0.1%) sterilized through a 0.22 µm pore filter into a sterile container and stored at 4 °C for up to 3 weeks ^23, 57^.

### Clonogenic and Spherogenesis assay in vitro

Single cell clonogenic assay was performed by depositing CD45^neg^/CD31^neg^/c-kit^pos^ and/or Sca-1^pos^ CSCs into 96-well gelatin-coated Terasaki plates by serial dilution. Individual CD45^neg^/CD31^neg^/c-kit^pos^ and/or Sca-1^pos^ cells were grown in CSC growth medium for 2 weeks when clones were identified and expanded. The clonogenicity of the CD45^neg^/CD31^neg^/c-kit^pos^ and/or Sca-1^pos^ cells was determined by counting the number of clones in each 96-well plate and expressed as a percentage of plated cells. A total of 5 plates were analyzed for each experiment. For cardiosphere generation, 1×10^5^ CD45^neg^/CD31^neg^/c-kit^pos^ and/or Sca-1^pos^ CSCs were placed in bacteriological dishes with CSC growth medium. Cardiospheres were counted per plate at 14 days and the number expressed as a percentage relative to the number of plated CSCs.

### Myogenic differentiation protocol in vitro

To perform in vitro myogenic differentiation of freshly isolated CSCs, immediately after isolation cells were plated onto laminin-coated 24-well plates or chamber slides to promote cell adhesion. Cells were maintained in a humidified incubator at 37°C with 5% CO₂ and allowed to adhere overnight in CSC growth medium. After attachment, the medium was replaced with a cardiac base differentiation medium, consisting of StemPro®-34 SFM differentiation medium (a serum-free medium conditioned with StemPro®-Nutrient Supplement, Gibco, Life Technologies), Ascorbic Acid (0.5 Mm, Sigma-Aldrich), 1-thioglycerol (4.5 × 10−4 M, Sigma-Aldrich), L-glutamine (2 mM), Non-Essential Amino Acids (Gibco, Life Technologies) and penicillin-streptomycin (1%, Life Technologies). For myocyte differentiation BMP4 (10ng/ml, Peprotech), Activin-A (50 ng/ml, Peprotech), β-FGF (10ng/ml, Peprotech), Wnt-11 (150ng/ml, R&D System) and Wnt-5a (150ng/ml, R&D System) for 4 days. At days 6 of cardiac differentiation, Dkk-1 (150ng/ml, R&D System) was added to base differentiation medium until day 14. Cell differentiation was evaluated at 14 days ^57^.

Alternatively, CSCs were placed in CSC growth medium in bacteriological dishes for 4 days for cardiospheres generation. Cardiospheres were then switched to base differentiation medium implemented with BMP-4 (10 ng/ml, Peprotech), Activin-A (50 ng/ml first day and then 10 ng/ml, Peprotech), β-FGF (10 ng/ml, Peprotech), Wnt-11 (150 ng/ml, R&D System) and Wnt-5a (150 ng/ml, R&D System). At day 8 cardiospheres were collected and transferred to laminin-coated dishes (1 µg/ml) and Dkk-1 (150 ng/ml, R&D System) was added to base differentiation medium. At day 14 differentiated cardiospheres were either trypsinized for RNA isolation or fixed with 4% PFA for immunohistochemistry.

### BrdU and EdU incorporation in vivo

To assess in vivo cell proliferation, twelve-sixteen weeks old double heterozygous TgMyh6-DTA mice underwent TAM or saline injection. Mice were then anaesthetized with isoflurane and implanted subcutaneously (between the scapulae) with osmotic minipumps (ALZET) to systemic release of Bromodeoxyuridine (BrdU, 50 mg/Kg/Day, Sigma-Aldrich) prepared by dissolving the powder in 50% deionized water and 50% DMSO ^26^. 28 days after TAM or Saline administration, mice were sacrificed, and hearts were fixed in formalin for immunohistochemistry analysis or enzymatically digested to obtain cardiac cell suspension. Additionally, a subset of twelve-sixteen weeks old double heterozygous TgMyh6-DTA mice were implanted following DTA-induced injury, with BrdU-releasing osmotic minipumps. From day 53 to 56 after TAM administration, mice received intraperitoneal injection of 5-Ethynyl-2’-deoxyuridine (EdU, 50 mg/Kg/Day, Life Technologies) for three consecutive days, dissolving the powders in 100 µl of 50% de-ionized water and 50% DMSO. Mice were then sacrificed, and hearts enzymatically digested to obtain cell suspension analyzed by flow cytometry (FACS). Cell-cycle entry was assessed by BrdU incorporation following intraperitoneal BrdU administration every 8 hours for two consecutive days before cell isolation, beginning immediately after TAM treatment.

### BrdU incorporation in vitro

Proliferation in CSCs clones was evaluated through BrdU incorporation assay. Single cell-derived cloned CSCs were plated at density of 0.5 × 10^4^ cells in 24 well dishes and initially maintained under proliferative conditions (growth medium). Cells were then switched to quiescence-inducing medium for 48h (DMEM F-12 implemented with 0.5% ESQ-FBS and LIF) to promote cell-cycle arrest. For reactivation, quiescent CSCs were re-exposed to mitogenic stimuli by replacing the quiescence-inducing medium with CSC growth medium and BrdU (10μM) was administrated every 8 hours. BrdU incorporation was measured after 48h of quiescence-inducing medium and after 24h of growth medium by immunostaining using the BrdU detection system kit (Roche). To assess proliferation on freshly isolated cells, BrdU (10μM) was administrated every 8 hours after 48 hours transfection with miR-221 precursor, anti-miR-221 or their relative controls. BrdU incorporation was then measured after 24 h by immunostaining using the BrdU detection system kit (Roche). Nuclei were stained with DAPI. Cells were evaluated using fluorescent microscopy.

### Transfection Reagent

Freshly isolated qCSCs and aCSCs were seeded at a density of 0.5 × 10^5^ cells in 12-well plates and incubate at 37 °C with 5% CO2 overnight in complete medium. The day after, qCSCs and aCSCs were transiently transfected with chemically modified oligonucleotides at a final concentration of 50 nM using Lipofectamine® RNAiMAX (Thermo Fisher Scientific) to over-express (pre miR-precursor) or inhibit (anti-miR inhibitor) miR-221 respectively (Ambion, Life Technologies) in OptiMem (Life Technologies). The mature miR-221 sequence used was the following: AGCUACAUUGUCUGCUGGGUUUC. A random miRNA sequence was used as pre-miR control or anti-miR control. Cells were incubated for 2–3 days at 37°C and then, analyze to assess transfection efficiency.

### Luciferase Assay

The 3′-UTR of mouse Cdkn1c (p57) containing the predicted miR-221 target sequences was cloned into the pEZX-MT06 dual luciferase vector (Genecopeia). qCSCs and aCSCs were transfected with pEZX-MT06 miR reporter vector for 48h. FLuc and RLuc activity was measured using the Luc-Pair Duo-Luciferase Assay Kit 2.0 according to the manufacturer’s instructions (Genecopeia).

### Quantitative RT-PCR (qRT-PCR)

RNA was extracted from clonogenic mouse CSCs using TRIzol Reagent (Ambion) and quantified using a Nanodrop 2000 Spectrophotometer (Thermo Fisher Scientific). Reverse transcription was performed with 0.5–1 µg of RNA using the High-Capacity cDNA Kit (Applied Biosystems). Quantitative RT-PCR was performed using TaqMan Primer/Probe sets (Applied Biosystems) using StepOne Plus real Time PCR System (Applied Biosystems). The following genes were tested: Cdk1, Cdk2, Cdc25a, Cdkn1a, Cdkn1b, Cdkn2a, Cdkn2b, Cdkn1c, Cdk6, Ccna2, Ccnb1, Ccnd1, Ccnd3, Ccnh, Cdc25c, E2f1, Mk67 as cell cycle genes, Cxcr4, Kdr, Bmi1, Ctnnb1, Hand1, Hand2, Mef2c, Gata4, Abcg2, Nkx2.5, Pou5f1, Tbx5, Wnt5a, Wt1 as stemness genes, Gata4, Nkx2.5, Mef2c, Hand2, Myh6, Myh7, Tnnt2, Actc1, Pln and Atp2a as myogenic genes, p57, Cre and miR221 (Supplementary Table 1). Data were processed by the ΔCt method using StepOne Software v2.3 and mRNA was normalized to the housekeeping gene, GAPDH. Heatmaps of qRT-PCR data were generated using color scales representing fold changes relative to the baseline value for each gene. All reactions were performed in technical triplicate.

### FACS analysis

Flow cytometry analysis was performed using FACSCanto II or LSRFortessa (BD Biosciences). FlowJo software (Treestar) was used to identify the percentage of cardiac cells expressing the different cell surface markers of interest. Appropriate labelled isotype controls were used to define the specific gates. Specific antibodies used are shown in Supplementary Table 2. To detect BrdU and EdU, cell suspensions were fixed and permeabilized. BrdU-labeled cells were detected using an anti-BrdU antibody (clone MoBU-1). EdU detection was performed using the EdU Click Proliferation Kit (BD) according to the manufacturer’s instructions. Freshly isolated CD45^neg^/CD31^neg^/c-kit^pos^ and/or Sca-1^pos^ cells were sorted using BD FACSAria III (BD Biosciences). For cell cycles profiles, cells were collected, washed twice with cold PBS, and fixed in 70% ethanol added dropwise while vortexing. Cells were stored at −20 °C for at least 2 h (or overnight) before analysis. Fixed cells were washed twice with PBS to remove ethanol and resuspended in FxCycle PI/RNAse Staining Solution (Life Technologies). Cells were incubated for 30 minutes at room temperature in the dark then analyzed using FACSCanto II or LSRFortessa (BD Biosciences). At least 10.000 events per sample were collected. Doublets were excluded by gating on FSC-A vs FSC-H. DNA content histograms were analyzed using software such as FlowJo. Cell cycle distribution (G0/G1, S, G2/M phases) was determined using the Watson or Dean-Jett-Fox model.

### Tissue Harvesting, Histology, and Immunohistochemistry

For immunohistochemistry analysis, cannulated hearts were arrested in diastole using cadmium chloride/potassium solution then fixed with 10% buffered formalin or with 4% PFA. The hearts were cut into apical, mid, and basal regions and right and left atria and embedded in paraffin or in Optimal Cutting Temperature Compound (OCT). Tissues were cut in 5 µm, 10 µm or 50 µm cross-sections. Formalin-fixed sections (5 µm and 10 µm) were stained with hematoxylin and eosin (H&E; Bio-Optica) according to standard protocols and with Masson’s trichrome stain (PolySciences, Inc.) for quantitative assessment of fibrosis. Cardiomyocyte cross-sectional area was measured through immunostaining with Wheat Germ Agglutinin Alexa Fluor 488 conjugate, WGA (1:200 dilution; Invitrogen) and digital analysis of acquired cardiac cross-section images (Leica, 1128 LAS AF Software). Cardiomyocyte diameter was measured across the nucleus on 3 transverse sections (∼500 myocytes/animal were sampled). For BrdU detection, antigen retrieval was achieved using Target Retrieval Solution, Citrate pH 6 (DAKO). Apoptotic cells were detected using a FITC-conjugated Anti-Terminal deoxynucleotidyl transferase kit (TdT, 1:50 dilution, In Situ Cell Death Detection Kit, Fluorescein, Life Technologies). For immunofluorescent staining, the following primary antibodies were used: anti-Cardiac Troponin I (1:100 dilution; Abcam), anti-BrdU (1:50 dilution, Sigma-Aldrich), anti-Ki67 (1:50 dilution, DAKO), anti-PCM1 (1:200, Atlas), anti-GFP (1:50, Rockland), anti-*α*-actinin. The specific primary antibodies were revealed by respective anti-rabbit IgG, anti-mouse or anti-goat IgG secondary antibody (1:100 dilution; Jackson Immunoresearch). The nuclei were counterstained with the DNA binding dye, 4, 6-diamidino-2-phenylindole (DAPI, Sigma-Aldrich) at 1 µg/mL. The number of necrotic/dead MF-20^pos^ cardiomyocytes was manually counted in cardiac cross sections for each power field using a 63× objective for a total of 20 fields, and the number of MF-20^pos^ cardiomyocytes was expressed as a percent fraction of the total cardiomyocyte number per mm^2^. The number of BrdU^pos^ cardiomyocytes was expressed as a percent fraction of the total cardiomyocyte nuclei. For mono/binucleation analyses, OCT embedded cardiac tissue was cut into 50μm sections. Sections were air dried for 10 minutes and antigen retrieval was achieved using Target Retrieval Solution, Citrate pH 6 (DAKO). Sections were blocked in 10% donkey serum. Tissue was then immunostained with the following primary antibodies: anti-mouse cardiac troponin (1:50), anti-rabbit PCM1 (1:100, Atlas), anti-BrdU (1:25, Abcam) and wheat germ agglutinin (WGA)-AlexaFluor555 (ThermoFisher) 3 hours or overnight on a shaker. Tissue was washed 3×10min with PBS and incubated in secondary antibodies, anti-mouse IgG, anti-rabbit IgG or anti-rat IgG respectively (1:100 dilution; Jackson Immunoresearch), for 3 hours on a shaker protected from light. Tissue was then washed 3×5mins in PBS and incubated in DAPI (1 µg/mL, Sigma-Aldrich). For analyses, each cardiomyocyte 3D cell boundary was determined in x, y and z using WGA, thus identifying the voxels (volumetric pixel) within each individual CMs. Unless otherwise specified, all staining were acquired and analyzed using confocal microscopy (LEICA STELLARIS SP5, Leica Microsystems).

### Cytospin

CD45^neg^/CD31^neg^/c-kit^pos^ and/or Sca-1^pos^ cardiac cells were suspended in complete CSC growth medium. A volume of 200 μl of a 5 × 10^4^ cells/ml suspension were directly loaded in each cytofunnel and spin down at 800 r.p.m. for 3 min onto poly-lysine-coated slides using a Shandon Cytospin 4 Cytocentrifuge (Thermo Fisher Scientific). Slides were immediately fixed using a spray fixative (BD Cytofix™ Fixation Buffer, BD Biosciences) and then processed for immunostaining as above described. Fluorescence was visualised and images acquired with confocal microscopy (LEICA Stellaris SP5).

### Statistical analysis

Statistical analysis was performed with GraphPad Prism version 9.00 for Windows (GraphPad Software 9). The quantitative data are reported as mean ± SD and binary data based on counts. The significance between the 2 groups was determined using Student’s t test. To draw comparisons between multiple groups, the ANOVA was used. A p value of < 0.05 was considered significant. Bonferroni’s post-hoc method was used to locate the differences. In these cases, the type 1 error (α = 0.05) was corrected by the number of statistical comparisons performed. For the in vitro cell and molecular biology experiments, giving the low number of the sample, the Kruskal–Wallis test (for multiple comparisons), and the Mann group –Whitney U test (for comparison between 2 groups) were performed.

## Results

### Cre-Inducible MYH6-restricted DTA activation acutely and selectively kills ventricular and atrial cardiomyocytes in adult transgenic mice

To assess the heart’s regenerative capacity after acute selective CM loss, we crossed transgenic B6129-Tg(Myh6-cre/Esr1)1Jmk/J mice (where the cardiomyocyte-specific alpha-myosin heavy chain - *Myh6*-promoter drives the expression of a Tamoxifen-inducible Cre recombinase fusion protein - mER-Cre-mER or MCM-, hereafter *Tg*Myh6^MCM^) ^53^ with homozygous Cre-reporter mice mutated in the ROSA26 locus to express a floxed-STOP sequence in front of the Diphtheria toxin A (DTA) gene (R26^stop-DTA^) ^58, 59^, in which tamoxifen (TAM) selectively triggers a Cre-mediated DTA expression in CMs, producing their acute death (Figure 1A). This *Tg*Myh6^MCM^ cardiomyocyte–specific, TAM-inducible line has been used in several landmark cell fate-mapping and functional gene analysis of adult CMs in mice ^53, 60, 61^.The more recently developed mice with Myh6^MCM^ knock-in alleles, despite their reported robust CM-restricted recombination, are not valid alternatives to the transgenic line because in these knock-in mice the targeted Myh6 allele is converted into a null, creating Myh6 haploinsufficiency ^62^. In double mutant TgMyh6^MCM^:R26^stop-DTA^ mice, hereafter *Tg*Myh6-DTA (n=3 male and n=3 females), Myh6 mRNA transcripts in atria and ventricles mirror wild-type controls, indicating physiological Myh6 expression levels (Supplementary Figure 1A,B). In contrast to Myh6, Cre mRNA was significantly higher in ventricular vs atrial CMs, independent of sex (Supplementary Figure 1C,D), suggesting greater Cre-recombination potential in ventricles.

**Figure 1.**
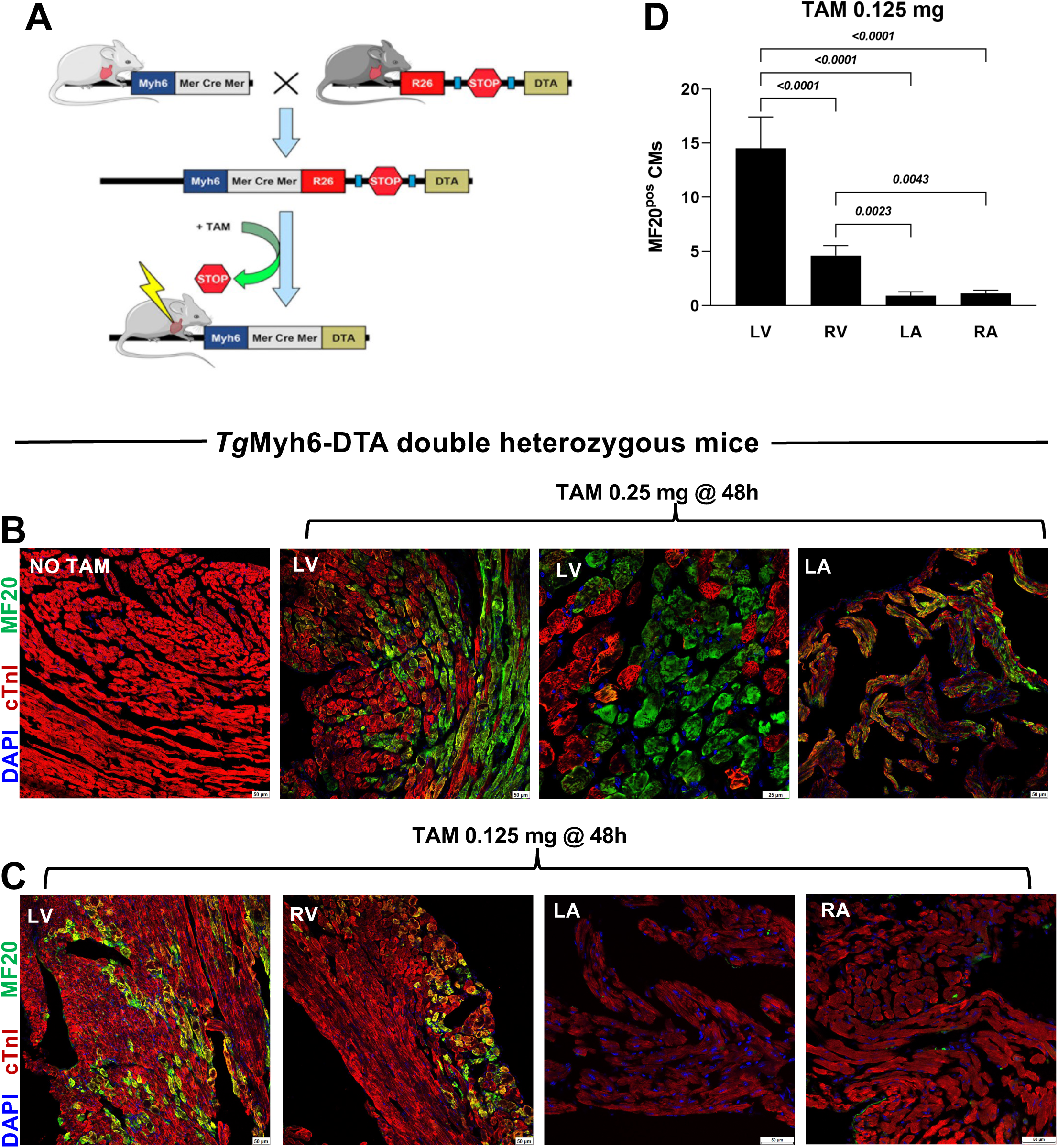
Dose dependent cardiomyocytes necrosis in *Tg*Myh6-DTA mice after TAM injection. (**A**) Schematic representation of double *Tg*Myh6-DTA mice generated by crossing a *Tg*-Myh6^MCM^ mouse expressing a tamoxifen (TAM)-inducible Cre recombinase fusion protein mER-Cre-mER (MCM) under the α-myosin heavy chain (Myh6) promoter, with a R26^stop-DTA^ reporter mouse that harbor a floxed STOP cassette upstream of the diphtheria toxin A (DTA) gene in the Rosa26 locus. Upon TAM administration, Cre-mediated recombination removes the STOP cassette, leading to DTA expression specifically in cardiomyocytes (CMs), resulting in their selective death. (**B-D**) Representative confocal images and bar graph showing necrotic CMs labelled *in vivo* with anti-myosin antibody (MF-20) in left and right ventricles as well as in left and right atria (LV, RV, LA, RA respectively) in *Tg*Myh6-DTA mice 48h after a single TAM doses of 0.25 mg (B) (Representative of n=1 heart sections) or 0.125 mg (C) (Representative of n=6 heart sections). (MF-20, green; cTnI, red; DAPI, blue nuclei. Scale bars=25 µm and 50 µm). All data are mean ± SD.

In trial-and-error experiments based on previous data^26, 63^, 0.5 mg TAM as a single dose IP in *Tg*Myh6-DTA mice, triggered within 2 days lethal DTA-mediated CM ablation in all 5 treated mice. The TAM dose halved to 0.25 mg still caused 80% mortality within two days (4/5 mice), with at least 65% necrosis of the LV’s CMs 2 days after TAM injection, as shown by *in vivo* MF-20 antibody labelling in the only surviving mouse (Figure 1B). Necrotic CMs displayed loss of membrane integrity, architectural disarray, and were mostly troponin (cTnI)-negative (due to troponin leakage), while overlapping MF-20/cTnI signals suggested early-stage necrosis (Figure 1B). To reduce mortality further, TAM was lowered to 0.125 mg, which resulted in ∼35% mortality (3/9 treated mice) within two days. This dose led to loss of 14.5± 2.9% and 4.6±0.9% of respectively left and right ventricular (LV and RV) CMs and 0.9±0,4% and 1.1±0.2% of respectively left and right atrial CMs, as confirmed and quantified by MF-20 labelling and necrotic features 48 hours after TAM injection (Figure 1C,D).

Thus, DTA activation in *Tg*Myh6-DTA mice selectively kills CMs in a dose dependent manner and it ablates more CMs in the ventricles than in the atria without detectably affecting the vascular and connective tissue cells. It should be emphasized that despite evident patchy areas of CM necrosis, the CM loss is diffuse throughout the myocardium, without any segmental myocardial loss.

### Complete functional cardiac recovery and cardiomyocyte renewal after selective cardiomyocyte loss by DTA activation

The significant CM loss by a single 0.125 mg dose of TAM in *Tg*Myh6-DTA mice poses a substantial biological and functional challenge, properly testing the adult heart’s intrinsic capacity for spontaneous repair from a pure CM loss with an intact blood supply and microvasculature. To investigate this response, fourteen *Tg*Myh6-DTA mice (8 males and 6 females) were injected as above. Four mice (3 males, 1 female) died within 24 hours post-TAM. Echocardiography of the remaining 10 mice 48h post-TAM, revealed increased left ventricular end-diastolic (LVEDD) and end-systolic (LVESD) diameters -- ∼10%, and ∼25% respectively -- compared to baseline (Figures 2A and Supplementary Table 3). These dilatations correlated with ejection fraction (EF) and fractional shortening (FS) depression (∼35% and ∼40%, respectively) (Figures 2A and Supplementary Table 3).

**Figure 2.**
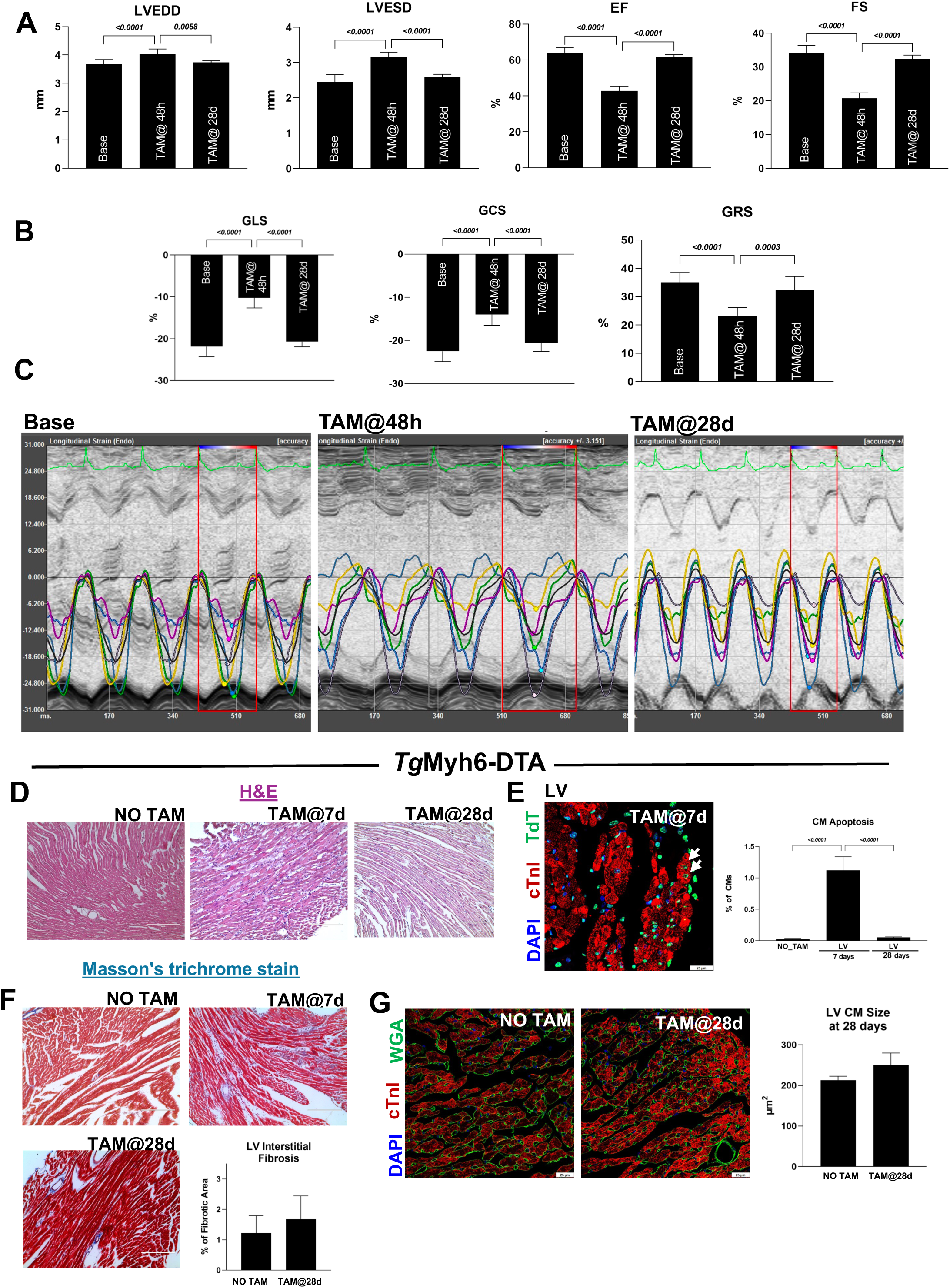
Cardiac function response to selective cardiomyocytes loss after DTA activation in *Tg*Myh6-DTA mice. (**A**) Bar graphs showing echocardiographic measurements of left ventricular end-diastolic (LVEDD) end-systolic (LVESD) diameters, ejection fraction (EF) and fractional shortening (FS) at baseline, 48h and 28 days after a single TAM dose of 0.125 mg in *Tg*Myh6-DTA mice. (**B,C**) Bar graph and representative images of regional speckle-tracking strain analysis showing the percentage of global longitudinal strain (GLS), global circumferential strain (GCS) and global radial strain (GRS) in *Tg*Myh6-DTA mice at baseline, 48h and 28 days after a single TAM at doses of 0.125 mg. (A-C) The number of animals per group is reported in the main text and in Supplementary Table 3. (**D**) Representative hematoxylin and eosin (H&E)-stained cross-sections of left ventricles from *Tg*Myh6-DTA mice at 7 and 28 days after TAM-induced injury, compared with saline-treated controls. Scale bar= 200 µm. (**E**) Representative confocal image and bar graph showing the percentage of apoptotic terminal deoxynucleotidyl transferase (TdT)-positive cardiomyocytes (CMs) (arrows) at 7 and 28 days after TAM-induced injury in *Tg*Myh6-DTA mice compared with saline-treated controls. (TdT, green; cTnI, red; DAPI, blue nuclei). Scale bar=25 µm. (**F**) Representative Masson’s Trichrome staining at 7 and 28 days after TAM-induced injury in *Tg*Myh6-DTA mice compared with saline-treated controls. Scale bars= 200 µm. Bar graph showing the quantification of interstitial fibrosis in *Tg*Myh6-DTA mice 28 days after TAM-induced injury compared with saline-treated controls. (**G**) Representative confocal images and bar graph showing CM size in *Tg*Myh6-DTA mice 28 days after TAM-induced injury compared with saline-treated controls (WGA, wheat germ agglutinin, green; cTnI, red; DAPI, blue nuclei. Scale bars= 25 µm). (D-G) Representative of n=5 heart sections from TAM injected *Tg*Myh6-DTA mice and n=3 heart sections from salina *Tg*Myh6-DTA mice. All data are mean ± SD

To further assess myocardial contractility and DTA’s effects on global and regional myocardial function, we performed speckle-tracking based strain analysis ^26, 54^. TAM decreased global longitudinal strain (GLS) by ∼50% at 2 days post-treatment (Figure 2B,C and Supplementary Table 3), with global circumferential strain (GCS) decreased by ∼40% (Figure 2B,C and Supplementary Table 3). Regional speckle-tracking strain analysis showed increased time to peak value and wall motion abnormalities at 2 days post-TAM (Figure 2C and Supplementary Table 3). Taken together, the functional parameters after TAM are consistent with and diagnostic of severe acute cardiac failure.

In agreement with the experiments represented in Fig. 1, seven days after administering 0.125 mg of TAM (n=5) or just saline (n=3) to *Tg*Myh6-DTA mice, extensive myocardial damage was confirmed (hereafter only LV data are presented, as the LV is the primary determinant of cardiac pump function and the chamber exhibiting the greatest acute necrosis in this model). In the LV, the CMs exhibited pronounced disarray together with an accumulation of inflammatory cells across the whole ventricular wall (Figure 2D and Supplementary Figure 2), which followed the extensive CM necrosis at two days (see Figure 1). At 7 days after DTA induction by TAM, 1.1±0.2% apoptotic CM nuclei, identified by Terminal Deoxynucleotidyl Transferase (TdT) immunostaining were detected as compared to 0.02±0.01% in saline injected control mice (Figure 2E).

Remarkably, despite the extensive cellular and functional damage produced, at 28 days post-TAM ventricular dimensions, EF and FS as well as global and regional strain values had returned to normal (Figures 2A-C and Supplementary Table 3).

Notably, the heart sections from *Tg*Myh6-DTA mice (n=5) at 28 days post-TAM showed a complete recovery in the global architecture of the heart as well as the individual CMs (Figure 2D and Supplementary Figure 2) together with a negligible number of apoptotic CMs (0.05±0.008%) (Figure 2E). Neither increased interstitial fibrosis (1.68± 0.8% in TAM- vs 1.22±0.6% in saline-injected *Tg*Myh6-DTA mice) (Figure 2F) nor evidence of CM hypertrophy (250.4±29.4 µm² in TAM- vs 212.8±9.7 µm² saline-injected mice) was detected at 28 days post-TAM (Figure 2G,H), documenting a robust cardiac repair that effectively compensated histologically and functionally for the acute loss of ∼15% of LV ventricular CMs.

To identify and quantify the newly-formed CMs, *Tg*Myh6-DTA mice (n=10) immediately after TAM injection were implanted subcutaneously (between the scapulae) with 28 days BrdU-releasing mini-osmotic pumps (n=7 surviving at 28 days)^25^. Additional *Tg*Myh6-DTA mice (n=7) were saline injected and similarly implanted with a BrdU minipump. Twenty-eight days post-TAM, heart cross sections from 4 mice per group were processed for BrdU immunohistochemistry, revealing a substantial number of newly formed BrdU-positive CM nuclei in TAM injected vs saline injected mice (8.05±0.9% vs 0.015±0.004%, respectively) (Figure 3A,B). Z-stack confocal microscopy shows that BrdU-positive CMs were exclusively mononucleated, in contrast to the mostly bi-nucleated surviving CMs (Figure 3C and Supplementary Figure 3). The remaining three hearts from each group were enzymatically digested to separate CMs and isolate CM nuclei. Flow cytometry of CM nuclei, dual-stained for PCM-1 ^2, 26, 64^ and BrdU, detected 2.2±0.8% BrdU-positive CM nuclei per heart in TAM vs 0.03±0.005% in saline injected mice (Figure 3D). This lower percentage of BrdU-positive nuclei measured by FACS in isolated CM nuclei compared with immunohistochemistry data likely reflects sensitivity differences between the assays. During CM dissociation and nuclear extraction, it is not possible to distinguish nuclei derived from mononucleated versus binucleated CMs (the latter comprise ∼80% of murine CMs) ^42, 65, 66^. In addition, substantial nuclei losses occur during CM and nuclear isolation. Accordingly, these FACS data should be interpreted just as qualitative confirmation of the tissue immunohistochemistry findings rather than as precise quantification.

**Figure 3.**
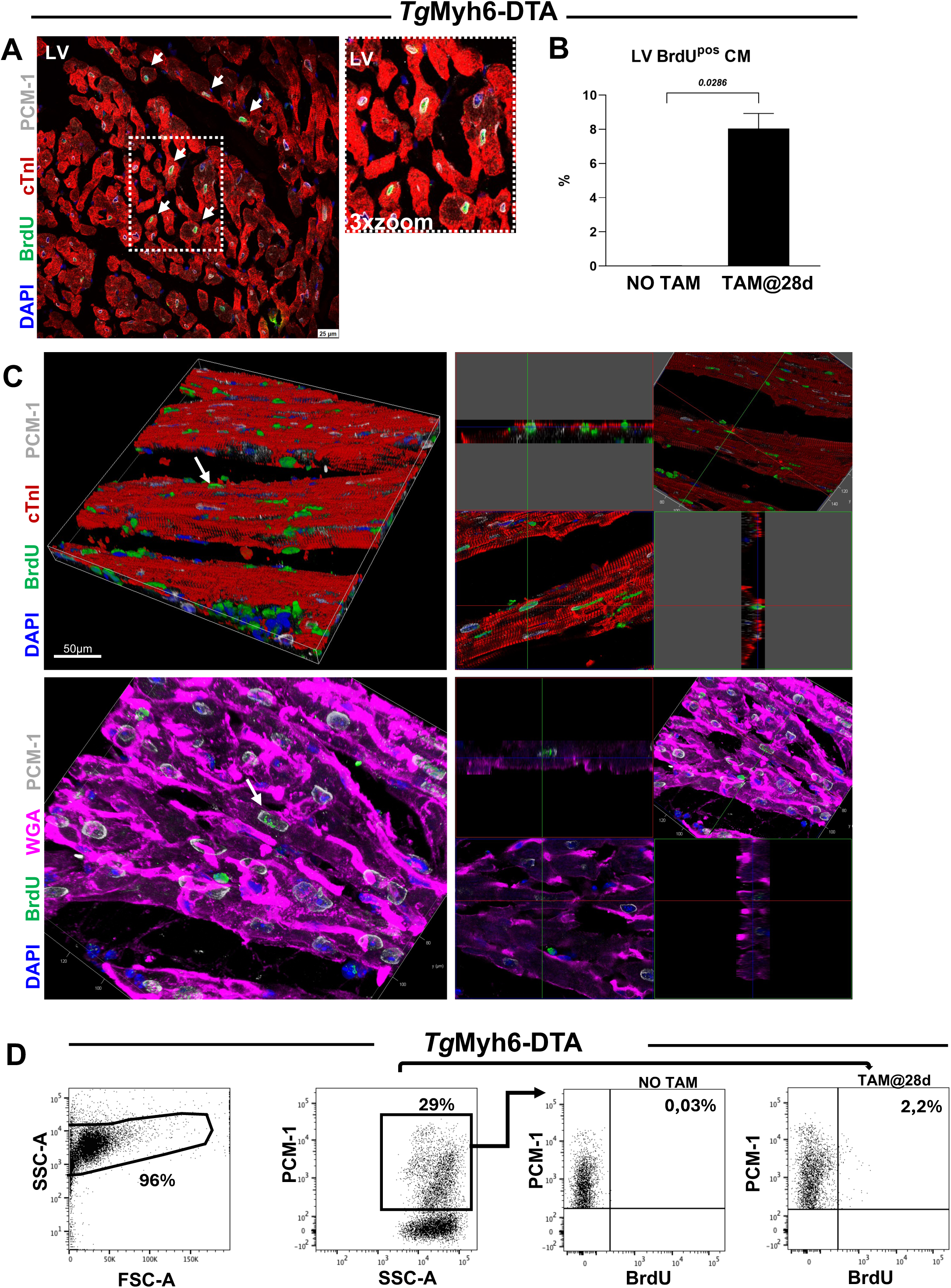
Regenerative capacity of the adult murine heart under selective CM ablation induced by DTA activation in *Tg*Myh6-DTA mice. (**A**,**B**) Representative confocal images and bar graph showing BrdU incorporation in left ventricular (LV) CMs (arrows) in TAM-induced injury *Tg*Myh6-DTA mice implanted with subcutaneous osmotic mini-pumps to systemically release BrdU for 28 days. Saline-treated *Tg*Myh6-DTA mice implanted with osmotic mini-pumps to systemically release BrdU for 28 days were used as controls (BrdU, green; cTnI, red; PCM-1, white; DAPI, blue nuclei). Scale bars = 25 µm. Representative of n=4 mice per group. **(C)** Representative 50 µm cardiac section and 3-dimensional (3D) volume of left ventricle showing mononucleated BrdU-positive CMs in TAM-induced injury *Tg*Myh6-DTA mice. Top panels: BrdU, green; PCM-1, white; cTnI, red; DAPI, blue nuclei. Bottom panels: WGA, magenta; DAPI, blue nuclei. Scale bars= 50 µm. Representative of n=3 per heart. (**D**) Representative flow-cytometry dot plots showing total cardiac nuclei identified by DAPI staining (left panel) and PCM-1-positive cardiomyocyte nuclei (middle-right panel) isolated from digested heart tissue of tamoxifen-induced *Tg*Myh6-DTA mice and saline-treated controls. Representative of n=3 biological replicates. All data are mean ± SD

Taken together, the foregoing findings reveal a robust CM regenerative capacity of the adult mammalian heart under selective diffuse (non-segmental) CM ablation. Indeed, the adult murine heart can replace up to ∼1/10 of its LV CM content in just one month. As expected from the histology, this surprisingly robust endogenous intrinsic regenerative response is accompanied by a complete functional recovery of the damaged heart.

### Cardiac stem cells, mostly quiescent in the adult heart, are activated to re-enter the cell cycle, regenerate the lost CMs and return to quiescence

To investigate the cellular origin of the new CMs generated after DTA-mediated injury, we first assessed the activation status of endogenous resident cardiac stem cells (CSCs) in adult *Tg*Myh6-DTA mice. In the rodent heart, CSCs have been identified within the CD45-negative/CD31-negative population expressing c-kit and/or Sca-1 ^15, 23, 55, 67^. However, because no single marker uniquely identifies bona fide CSCs, their direct isolation and quantitative assessment in vivo remain challenging. Clonogenic, spherogenic, and differentiation in vitro assays have shown that approximately 20% of freshly isolated CD45-negative/CD31-negative/c-kit-positive/Sca-1-positive cells display CSC properties; however, this proportion is likely a gross underestimate because of the limited efficiency of these functional assays ^15, 23, 55, 67^.

Previously, we isolated CSCs as CD45-negative/CD31-negative/c-kit-positive cells, approximately 50–60% of which co-expressed Sca-1 at isolation and became uniformly Sca-1-positive after culture expansion^15, 23, 55, 67^. In the present study, because several reports have also isolated multipotent cardiac stem/progenitor cells using Sca-1 as the principal marker,^15,16,19,22^ we adopted a broader, marker-inclusive strategy. Following CD45/CD31 depletion, cells positive for c-kit and/or Sca-1 were isolated by FACS to maximize recovery of the resident CSC compartment without restricting its definition to a single marker (Figure 4A). The resulting freshly isolated population comprised approximately 20% c-kit-positive cells, about half of which co-expressed Sca-1, and approximately 80% Sca-1-positive cells, the large majority of which were c-kit-negative (Figure 4A).

**Figure 4.**
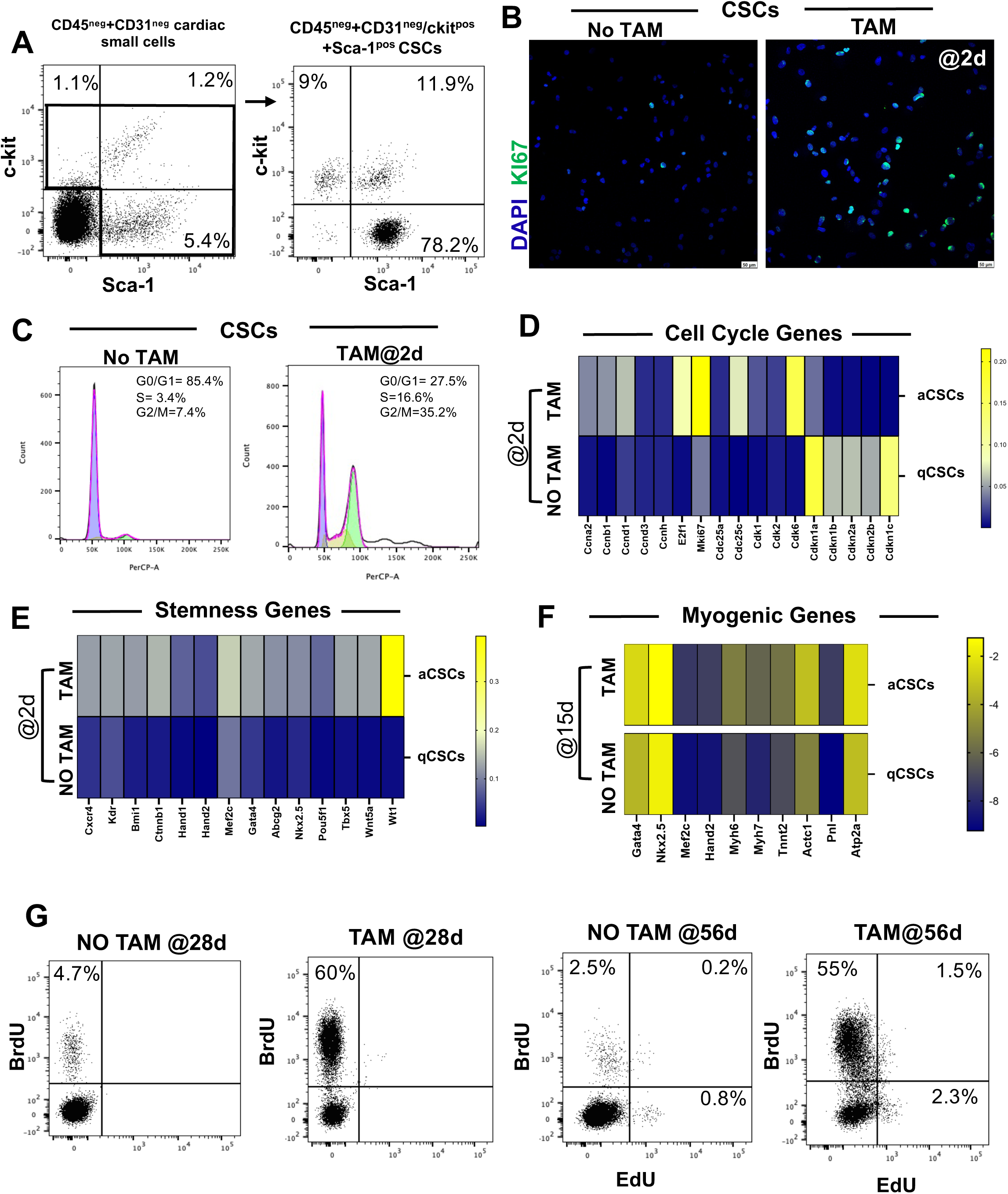
Endogenous cardiac stem cell activation after selective CM ablation induced by DTA activation in *Tg*Myh6-DTA mice. (**A**) Dot plots showing the percentage of c-kit^pos^ and Sca-1^pos^ cardiac stem cells (CSCs) after CD45 and CD31-negative MACS sorting (left panel). Right panel show the purity of c-kit^pos^ and Sca-1^pos^ FACS sorted CSCs. Representative of n=4 biological replicates. (**B**) Representative confocal images from a cytospin preparation showing the percentage of Ki67^pos^ positive CD45^neg^/CD31^neg^/c-kit^pos^/Sca-1^pos^ CSCs in *Tg*Myh6-DTA mice 2 days after saline or TAM-induced injury (Ki67, green; DAPI, blue nuclei). Scale bar=50 µm. Representative of n=4 biological replicates. (**C**) Representative flow cytometry cell-cycle histograms of CSCs from *Tg*Myh6-DTA mice 2 days after saline or TAM-induced injury. The first and last peaks correspond to the G0/G1 and G2/M phases, respectively, and the intermediate region corresponds to the S phase. Representative of n=4 biological replicates. (**D**, **E**) Heatmaps showing qRT-PCR analysis of genes involved in cell cycle progression (D) and stemness (E) in CSCs from *Tg*Myh6-DTA mice 2 days after saline (qCSCs) or TAM-induced injury (aCSCs). (**F**) Heatmap showing qRT-PCR analysis of the main myogenic genes in CSCs from *Tg*Myh6-DTA mice 15 days after saline (qCSCs) or TAM-induced injury (aCSCs). (D-F) The color scale represents changes in Ct values (threshold cycle) relative to the GAPDH-normalized control. Representative of n=3 biological replicates per group, each performed in technical triplicate. (**G**) Flow cytometry dot plots showing BrdU incorporation in CSCs from *Tg*Myh6-DTA mice 28 days after saline or TAM-induced injury (left and middle panels) and BrdU and EdU labelling after 56 days (right panels). Representative of n=3 biological replicates. All data are mean ± SD

In control saline-injected *Tg*Myh6-DTA hearts (n=4), most CD45^neg^/CD31^neg^/c-kit^pos^ and/or Sca-1^pos^ cells ^23, 57, 68^(hereafter just CSCs for the sake of brevity) were quiescent, with only 3.4 ±0.9% expressing the proliferation marker Ki67 (Figure 4B). In contrast, two days after TAM injection (n=4) a robust CSC activation was evident, with 47.3 ± 7% of the CSCs having exited quiescence and entered the cell cycle as shown by Ki67 labelling (Figure 4B). Consistent with these findings, cell-cycle profiling by FACS showed that CSCs from control hearts were almost entirely in G0/G1 (85.8±1.3%), with only a small fraction in S phase (7.6±1.15%) and virtually no cells in G2/M (1.13±0.5%). Whereas, CSCs isolated 2 days after DTA-induced injury displayed a marked redistribution across the cell cycle, with a reduction in the G0/G1 fraction (34.1±6.1%) and a concomitant increase in S-phase (16.1±1.5%) and G2/M-phase cells (31.4±3.4%), indicating active cell-cycle progression (Figure 4C). Because only a subset of CD45^neg^/ CD31^neg^ /c-kit^pos^/Sca-1^pos^ are true CSCs ^23, 55^, it is likely that the cells which have remained quiescent during the severe DTA-injury challenge are either not CSCs or not within the proper microenvironment for activation.

To identify the molecular underpinnings of this response, we performed multigene qRT-PCR on CSCs isolated 2 days after control (saline-injected, n=3) and TAM-treated (n=3) *Tg*Myh6-DTA hearts, respectively quiescent CSCs (qCSCs) and activated CSCs (aCSCs). aCSCs showed increased expression of genes related to cell-cycle activation, proliferation and mitosis (Ccna2, Ccnb1, Ccnd1, Ccnd3, Ccnh, E2f1, Mk67/Ki67, Cdc25a, Cdc25c, Cdk1, Cdk2, and Cdk6) and to cell stem/progenitor phenotype (Cxcr4, Kdr, Bmi-1, Ctnnb1, Hand1, Hand2, Mef2c, Gata4, Abcg2, Nkx2.5, Pou5f1, Tbx5, Wnt5A, and Wnt1) (Figure 4D,E). In contrast, the expression of genes acting as cyclin-dependent kinase inhibitors (Cdkn1a/p21^Cip1-Waf1^, Cdkn1b/p27^Kip1^, Cdkn2a/p16^INK4^, Cdkn2b/p15^INK4b^, Cdkn1c/p57^Kip2^) was reduced in aCSCs 2 days after TAM administration compared to saline qCSCs (Figure 4D). At 15 days post-saline (n=3)or TAM injections (n=3), qRT-PCR for selected myogenic genes revealed consistent cardiac transcription factors and cardiac contractility genes upregulation (Gata4, Nkx2.5, Mef2C, Hand2, Myh6, Myh7, Tnnt2, Actc1, Pnl, and Atp2A) in aCSCs indicating injury-activation of a cardiomyogenic differentiation program (Figure 4F).

To determine whether activated CSCs return to a quiescent state *in vivo* once the damage has been corrected, we assessed CSC proliferation using a BrdU-releasing osmotic pump for 28 days after DTA-induced damage in *Tg*Myh6-DTA mice (n=6), which were then administered EdU intraperitoneally for 3 consecutive days, after fully recovered cardiac function (i.e. from day 53 to 56 after TAM). At 28d post TAM, 60±10% of the CSC cohort had entered the cell cycle and were BrdU positive (Figure 4G). In contrast, EdU labelling for 3 days at 8 weeks post-DTA revealed that only a small fraction of CSCs (3.8±1%) was cycling, a value comparable to uninjured controls (Figure 4G). Notably, ∼40% of EdU-positive CSCs were also BrdU-positive (Figure 4G), while >90% of the BrdU-positive were EdU negative, indicating that most cells activated by DTA injury (BrdU-positive) had subsequently returned to quiescence, thus completing a full activation-return to-quiescence cycle characteristic of regenerative progenitor cells that return a quiescent state after accomplishing tissue repair ^69–75^. These data show that a small cohort of these cells remained in the cell cycle long after the primary injury, possibly to support the low steady-state CM turnover in the adult heart.

### Cardiac stem cells are the main effectors of cardiomyocyte regeneration after severe diffuse myocardial damage

Having established that endogenous CSCs exit quiescence and enter the cell cycle, their progeny initiates a cardiomyogenic program and they return to quiescence after regenerating the DTA-induced CM loss, we next asked whether activated CSCs are required for myocardial regeneration and, if so, whether they are sufficient to repair the damage. To this end, we used a previously validated low-dose 5-fluorouracil (5-FU) to ablate the injury-activated proliferative cardiac compartment which includes the CSCs ^25, 26, 76^. Because 5-FU targets cycling cells irrespective of lineage, this approach suppresses proliferating CSCs as well as any other cell type that may re-enter the cell cycle after injury, including, in principle, rare adult terminally differentiated CMs re-entering the cell cycle. Thus, this strategy allows to determine whether regeneration depends on cell proliferation, but not to assign that requirement to a specific cell type.

Two days after TAM injection, *Tg*Myh6-DTA mice (n=16) were implanted with subcutaneous osmotic minipumps to deliver 5-FU (n=10) or just saline (n=6) continuously over 25 days (Figure 5A). Importantly, we have previously established that the dose and administration regimen of 5-FU used in the present study are not intrinsically cardiotoxic and do not affect cardiac tissue structure or function *per se.* ^25, 26^ 5-FU–treated *Tg*Myh6-DTA mice surviving at 28 days post-TAM (n=5) showed significant ventricular dilation and systolic dysfunction compared to FU-untreated mice (saline injected, n=5). (Figure 5B,C, Supplementary Figure 4 and Supplementary Table 4). Histological analysis of 5-FU–treated hearts revealed a severely reduced CSC number (0.02±0.008/mm² vs 0.06±0.015/mm² in saline injected mice) (Figure 5D) and practically no Ki67^pos^ CMs (0.002±0.001% vs 0.82±0.35%) (Figure 5E,F), confirming the failure of regenerative response upon ablation of proliferating cells which resulted in pronounced hypertrophy of surviving CMs (358.7±27.9 µm² vs 199.26±19.8 µm² in saline injected mice) (Figure 5G) and apoptosis (0.9±0.2% vs 0.05±0.03%) (Figure 5H).

**Figure 5.**
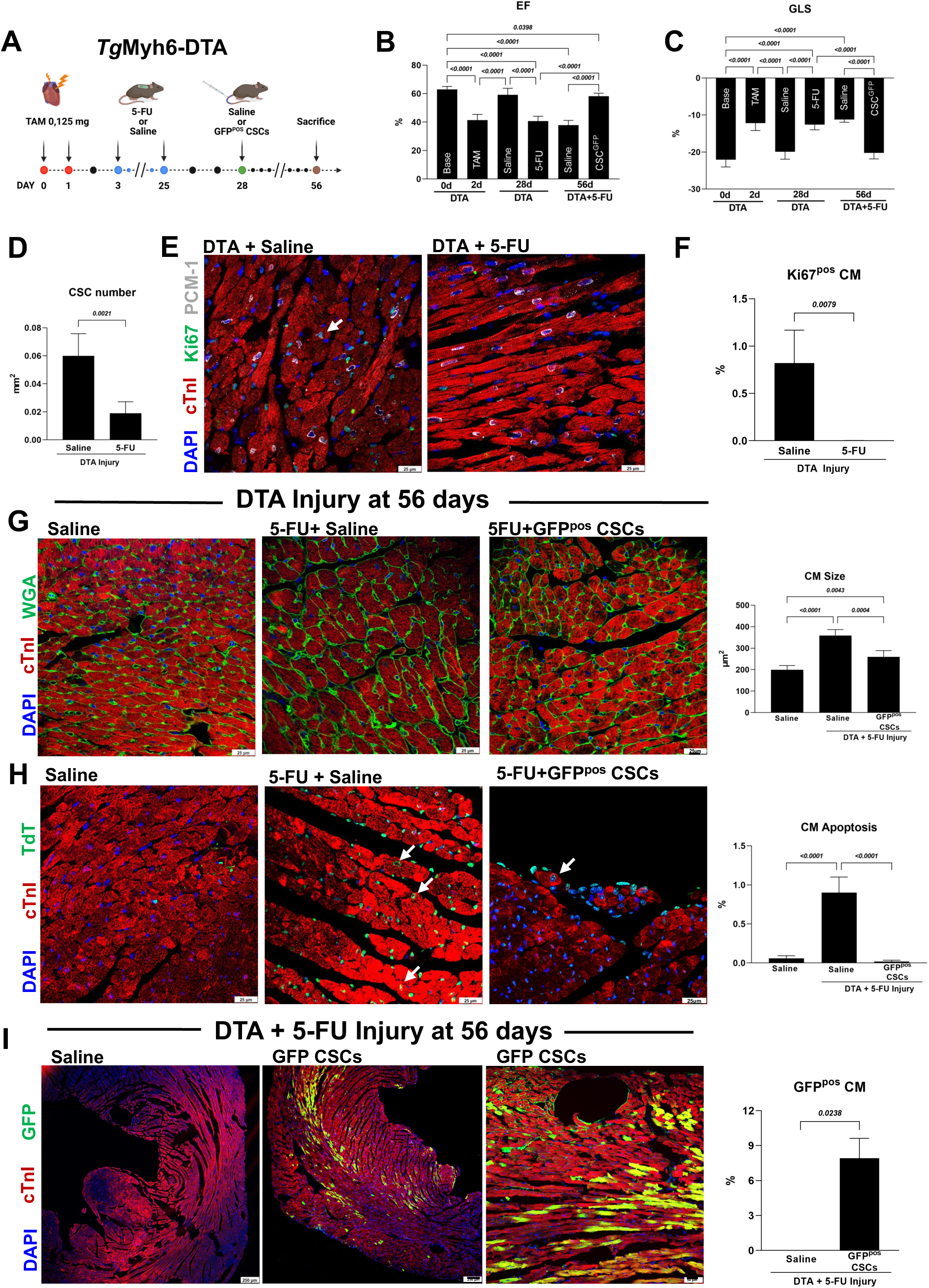
DTA-Induced diffuse damage triggers CSC activation, proliferation, and differentiation into mature CMs. (**A**) Experimental design of the in vivo 5-FU-based approach to ablate proliferating cardiac cells after TAM-induced injury in *Tg*Myh6-DTA mice with or without intravenous transplantation of GFP-labeled clonogenic CSCs. (**B**,**C**) Bar graphs showing ejection fraction (EF) and global longitudinal strain (GLS) at baseline, 2 days after TAM-induced injury, 28 days after TAM-induced injury in saline- or 5-FU-treated *Tg*Myh6-DTA mice and 56 days after injury in saline- or 5-FU+CSCs^GFP^-treated *Tg*Myh6-DTA mice. The number of animals used in each experimental group is reported in the main text and in the relative Supplementary Table 4. (**D**) Bar graph showing the number of CSCs per mm^2^ in heart sections collected 28 days after TAM-induced injury in saline or 5-FU-treated *Tg*Myh6-DTA mice. Representative of n=5 biological replicates. (**E**) Representative confocal images showing the percentage of Ki67^pos^/PCM-1^pos^ CMs 28 days after TAM-induced injury in saline or 5-FU-treated *Tg*Myh6-DTA mice. Representative of n=5 biological replicates. (Ki67, green; cTnI, red; PCM-1, white; DAPI, blue nuclei, scale bars= 25 µm. (**F**) Representative bar graph showing the percentage of Ki67^pos^ CMs 28 days after TAM-induced injury in saline or 5-FU-treated *Tg*Myh6-DTA mice. Representative of n=5 biological replicates. (**G**) Representative confocal images and bar graph showing CM size measurement 56 days after TAM-induced injury in saline, 5-FU+saline and 5-FU+ CSCs^GFP^ - treated *Tg*Myh6-DTA mice. (WGA, green; cTnI, red; DAPI, blue nuclei, scale bars=25 µm. (**H**) Representative confocal images and bar graph showing the percentage of apoptotic terminal deoxynucleotidyl transferase (TdT)-positive CMs in TAM-induced injury in saline, 5-FU+saline and 5-FU+ CSCs^GFP^ -treated *Tg*Myh6-DTA mice. (TdT, green; cTnI, red; DAPI, blue nuclei, scale bar= 25 µm). (**I**) Representative confocal images and bar graph showing the percentage of newly generated GFP^pos^ CMs in TAM-induced injury in saline, 5-FU+saline and 5-FU+ CSCs^GFP^ -treated *Tg*Myh6-DTA mice. (GFP, green; cTnI, red; DAPI, blue nuclei, Scale bars = 250 µm and 50 µm). (G-I) Representative of n=6, n=3 and n=6 biological replicates for saline, 5-FU+saline and 5-FU+ CSCs^GFP^ treated *Tg*Myh6-DTA mice respectively. All data are mean ± SD

To test whether CSCs could rescue this proliferative and regenerative defect, additional 5-FU–treated *Tg*Myh6-DTA mice were randomized to receive intravenous injection of either saline (n=6) or 4×10^5^ GFP-labelled clonogenic mouse syngeneic CSCs (CSCs^GFP^ n=6)^25, 26, 76^ (Figure 5A). The transplanted CSCs are derived from a single-cell CSC clone which maintains its myogenic character ^57^. Twenty-eight days after injection, all CSCs^GFP^-treated mice (6/6) had survived and fully recovered cardiac function whereas only 3 out of 6 saline-injected animals survived but remained in overt cardiac failure (Figure 5B,C, Supplementary Figure 4 and Supplementary Table 4). CSCs^GFP^-treated mice as compared to saline-injected mice show a reduced CM size (259.07±29.56 µm² in 5-FU+CSCs^GFP^ vs 358.7±27.9 µm² in 5-FU+saline) (Figure 5G), and reduced CM apoptosis (0.018±0.017% in 5-FU+CSCs^GFP^ vs 0.9±0.2% in 5-FU+saline) (Figure 5H). Importantly, the CSCs^GFP^ adoptive transfer resulted in the formation of 7.9±1.7% GFP-positive CMs, with many new CMs organized in islands suggestive of clonal expansion (Figure 5I). Therefore, clonal expanded CSCs behave as true adult stem/progenitor cells in the *in vitro* and the *in vivo* myocardial regeneration essays shown here, in agreement with previous results ^12, 25, 55^.

Taken together, these results demonstrate that CSCs are the main effectors of CM renewal and for anatomical and physiological cardiac repair following extensive diffuse myocardial injury.

Additionally, these data identify CSCs as the principal source of the new CMs in this model, whereas a contribution from proliferation of pre-existing CMs, if present, would not be detected under our experimental conditions. However, the magnitude of new CM formation shown here is many-folds higher than by the most effective replication of mature differentiated CM protocol reported so far ^41–44^. Collectively, the results shown above demonstrate that selective but diffuse CM loss by DTA triggers enough CSC activation, proliferation, and differentiation into mature CMs to repair the damage, followed by their return to the quiescent state.

### Injury-induced exit from the quiescent state fosters CSCs regenerative potential

Although the adoptive-transfer experiments showed that the progeny of a single CSC is sufficient to rescue myocardial regeneration, this clone had been selected for its robust myogenicity and it had undergone expansion in vitro for many generations ^57^, therefore, it did not directly address the functional consequences of physiological CSC activation within the injured heart. To determine more directly whether acute injury-induced exit from quiescence enhances the regenerative potential of endogenous resident CSCs *in situ*, we isolated qCSCs from saline-treated TgMyh6-DTA mice (n=6) and injury-aCSCs from TgMyh6-DTA (n=6) mice 2 days after TAM-induced myocardial damage, then compared their proliferative, clonogenic, cardiomyogenic, and regenerative properties *in vitro* and *in vivo*. Cell cycle entry was assessed by BrdU incorporation following in vivo ip BrdU administration every 8 hours for 2 consecutive days before cell isolation, starting immediately after TAM treatment.

After isolation, cells were used for *in vitro* functional assays. The number of BrdU-incorporating cells in 2 days was significantly higher in aCSCs (60 ± 6.8%) than in qCSCs (5.5 ± 1.1%) (Figure 6A).

**Figure 6.**
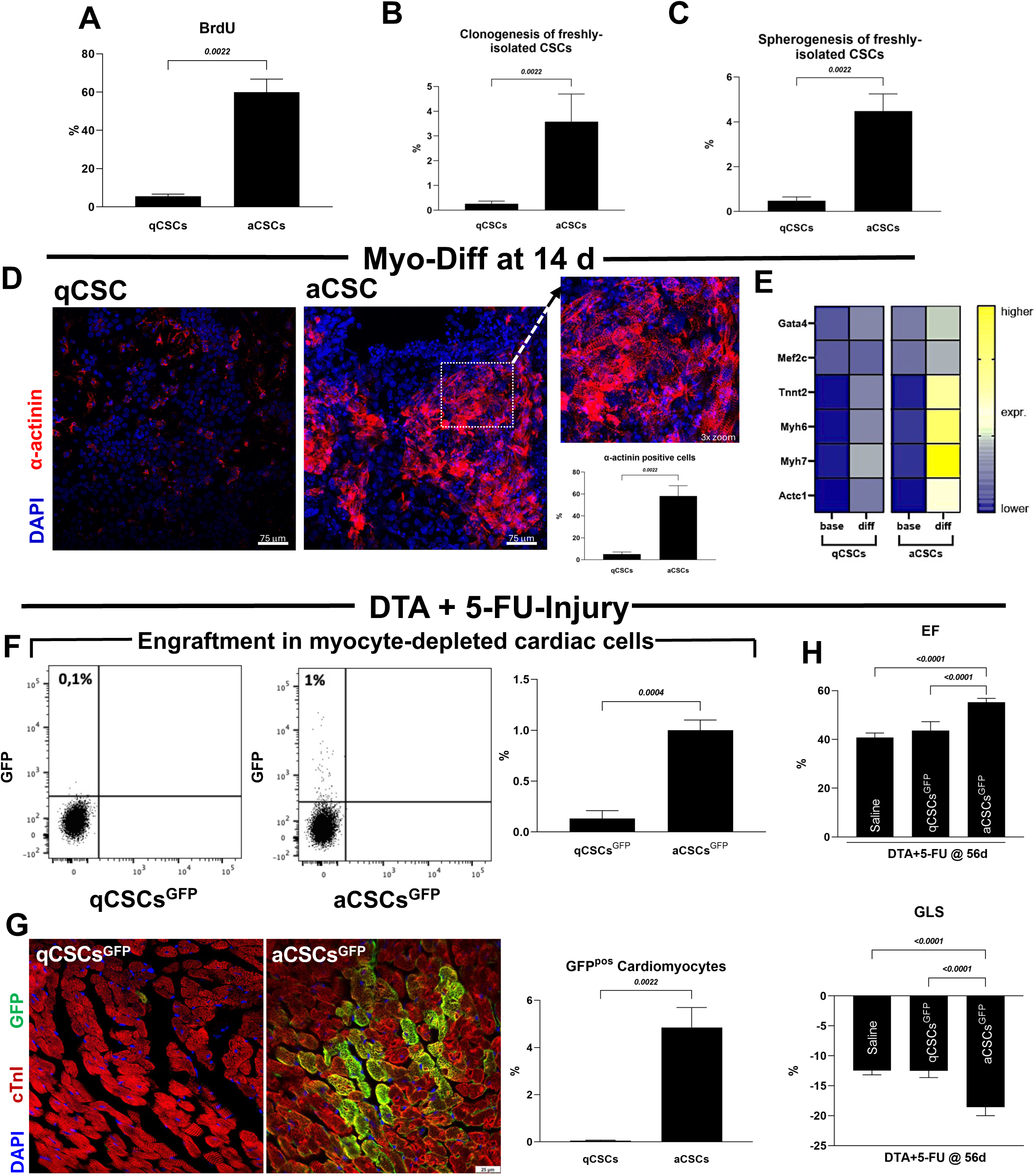
DTA-Induced diffuse CM loss drives CSC exit from quiescence inducing cardiac regeneration. (**A**) Bar graph showing the in vitro percentage of BrdU incorporation in freshly isolated quiescence CSCs (qCSCs) and activated CSCs (aCSCs) isolated from *Tg*Myh6-DTA mice 2 days after salina or TAM-induced damage. Representative of n=6 biological replicates, each performed in technical triplicate. (**B**, **C**) Bar graphs showing the in vitro percentage of clones and cardiospheres obtained from freshly isolated qCSCs and aCSCs from *Tg*Myh6-DTA mice 2 days after salina or TAM-induced damage. Representative of n=6 biological replicates, each performed in technical triplicate. (**D**,**E**) Representative confocal images, bar graph and heatmap showing enhanced cardiomyogenic potential in aCSCs compared to qCSCs isolated from *Tg*Myh6-DTA mice 2 days after salina or TAM-induced damage and cultured in vitro for 14 days in pro-differentiation medium (actinin, red; DAPI, blue nuclei, scale bars=75 µm). The heat map color scale represents changes in Ct values (threshold cycle) relative to the GAPDH-normalized control. Representative of n=6 biological replicates. (**F**) Representative dot plots and bar graph showing the percentage of engrafted qCSCs^GFP^ and aCSCs^GFP^ obtained from *Tg-Myh6^MCM^:R26^eGFPstop-DTA^*donor mice two days after saline or TAM-induced damage respectively and intravenously injected into 5-FU–treated *Tg*Myh6-DTA mice 28 days post-TAM. Representative of n=3 biological replicates. (**G**) Representative confocal images and bar graph showing the percentage of GFP^pos^ CMs at 56 days in 5-FU–treated *Tg*Myh6-DTA mice, 28 days after TAM or salina administration of qCSCs^GFP^ and aCSCs^GFP^(GFP, green; cTnI, red; DAPI, blue nuclei, scale bars = 25 µm). Representative of n=6 replicates. (**H**) Bar graphs showing the percentage of the ejection fraction (EF) and the global longitudinal strain (GLS) in 5-FU–treated *Tg*Myh6-DTA mice 28 days post-TAM or salina and injected with qCSCs^GFP^ and aCSCs^GFP^ from *Tg-Myh6^MCM^:R26^eGFPstop-DTA^* donor mice two days after saline or TAM-induced damage respectively. The number of animals used in each experimental group is reported in Supplementary Table 5. All data are mean ± SD

Clonogenic assays showed that freshly isolated qCSCs had minimal colony-forming capacity (0.3± 0.17%), while 3.6 ± 1.1% of aCSCs display clonal formation (Figure 6B). Similarly, when plated on bacteriological dishes, freshly isolated aCSCs exhibited significantly greater spherogenicity than qCSCs (4.5± 0.8% vs 0.5± 0.2%, respectively) (Figures 6C).

To determine the CSCs cardiomyogenic potential, we subjected both quiescent and activated CSCs to a standardized *in vitro* differentiation protocol. After 14 days in pro-differentiation medium, aCSCs yielded a significantly higher number of cells expressing α-actinin (58.1± 9.3% vs 5.4±3.6% in activated and quiescent CSCs, respectively), with organized sarcomeric structures clearly visible by immunofluorescence (Figure 6D) and confirmed by RT-PCR analysis of several CM-specific genes (Figure 6E), which established that CM-selective injury not only promotes the expansion but also primes the CSCs toward cardiomyogenic commitment.

To test whether the quiescent-to-active transition impacts CSC regenerative capacity *in vivo*, we isolated GFP-labelled CSCs from *Tg-Myh6^MCM^:R26^eGFPstop-DTA^* donor mice ^77^ two days after saline (n=10) or TAM treatment (n=6) (respectively, quiescent and activated CSCs^GFP^, heretofore qCSCs^GFP^ and aCSCs^GFP^) and intravenously (i.v.) injected them (4×10^5^ cells) into 5-FU–treated *Tg*Myh6-DTA mice 28 days post-TAM. At 28 days post-cell adoptive transfer (see Methods and schematic representation in Figure 5A), aCSCs^GFP^ had an increased engraftment. Indeed, GFP^pos^ cells within myocardial LV tissue rose from 0.1±0.07% after qCSCs^GFP^ to 1±0.1% after aCSCs^GFP^ (Figure 6F). Importantly, injury-induced aCSCs significantly boosted new CM formation as revealed by GFP-positive CMs (0.05 ± 0.02% after qCSCs^GFP^ and 4.9± 0.8% after aCSCs^GFP^ transfer) (Figure 6G) confirming homing, differentiation and tissue integration. Functionally, only aCSCs^GFP^ improved cardiac function with significantly enhanced ejection fraction and global strain (Figure 6H, Supplementary Figure 5 and Supplementary Table 5). In contrast, qCSCs^GFP^ produced no functional recovery (Figure 6H, Supplementary Figure 5 and Supplementary Table 5). The high myogenicity of the aCSCs^GFP^ which is not very different from that obtained with the progeny of a myogenic clone, further support the notion that most of the CD45^neg^/CD31^neg^/c-kit^pos^/Sca-1^pos^ are true CSCs.

Taken together, these findings strongly support the conclusion that injury-activated CSCs are the source of the cardiomyocytes regenerated after DTA-induced injury. Furthermore, they show that the aCSCs injected i.v. home to the damaged myocardium in large enough numbers to induce regeneration of the damaged myocardium.

### miR-221 regulates CSC transition from quiescence to activation via p57 repression following DTA-induced myocardial injury and *vice versa* after repair

To gain insight into the molecular mechanisms governing CSC transition from quiescence to activation, we focused on p57 (Kip2), a cyclin-dependent kinase inhibitor, and on miR-221 target that is a well-established enforcer of quiescence in multiple adult stem cell compartments ^78–82^. qRT–PCR showed that p57 was downregulated ∼35-fold in aCSCs isolated from DTA-injured hearts 48 hours after TAM (n=3) compared with qCSC controls (n=3) (Figure 7A). In parallel, miR-221, which is highly expressed in clonogenic CSCs ^57^ and is known to target both p57 and p27 in other cellular contexts ^80^, was upregulated ∼40-fold in aCSCs from DTA-injured hearts (Figure 7B). Together with the observation that p27 is also downregulated in activated CSCs after DTA-induced injury (Figure 4), these findings suggest that miR-221 may promote CSC activation through coordinated repression of Cip/Kip family members. Nevertheless, because p57 is more strongly linked to stem cell quiescence control, we prioritized it for further mechanistic analysis.

**Figure 7.**
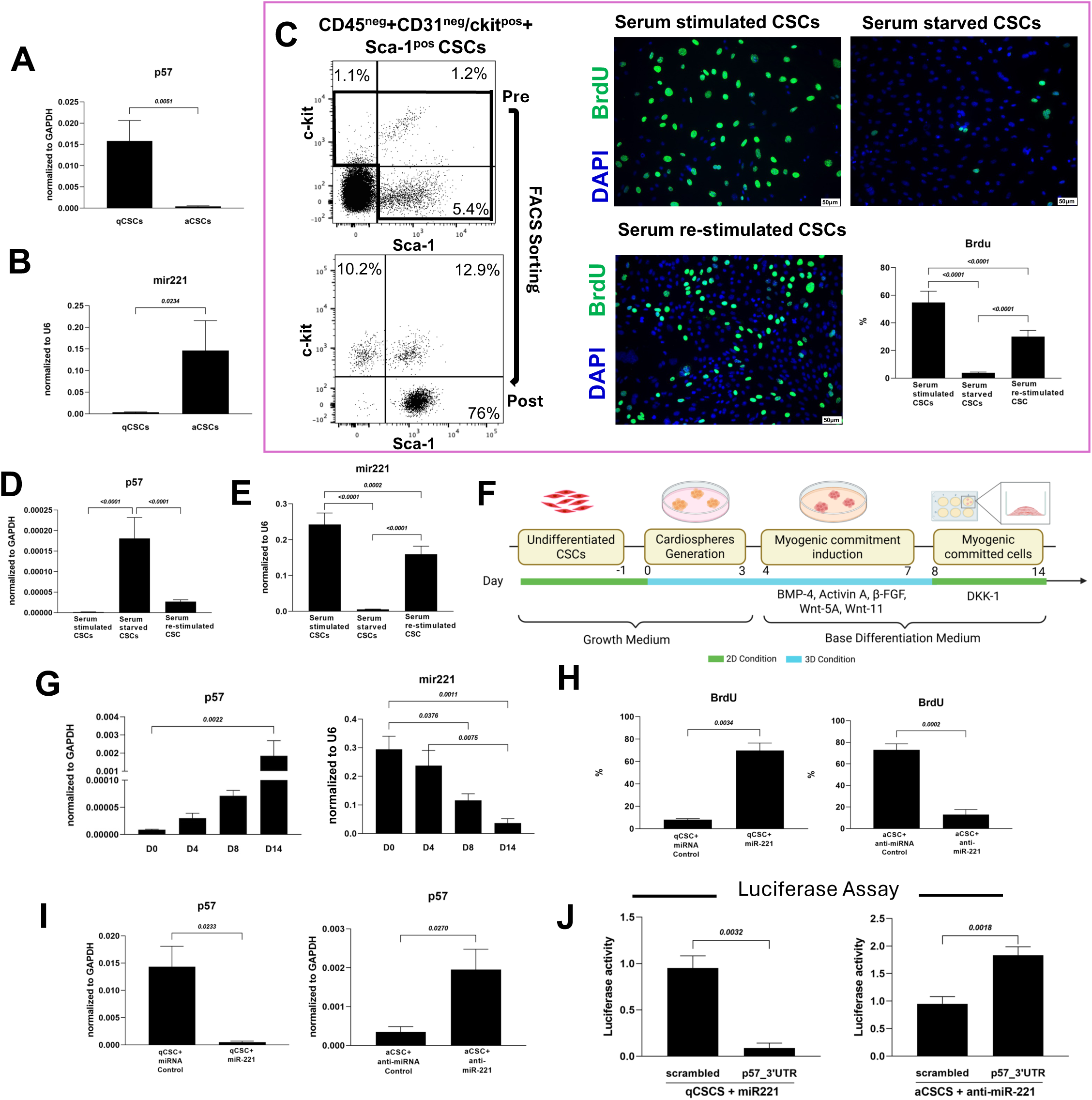
miR-221/p57 axis regulates CSC quiescence-to-activation transition. (**A,B**) qRT-PCR analysis showing the expression of p57 and miR221 in qCSCs and aCSCs from *Tg*Myh6-DTA mice 2 days after saline or TAM-induced damage. (**C**) Representative dot plots, confocal images, and bar graphs showing BrdU incorporation in clonogenic CSCs maintained under proliferative conditions, cultured in quiescence-inducing medium, or induced to re-enter the cell cycle by re-exposure to mitogenic stimuli. (BrdU, green; DAPI, blue nuclei, scale bars=50 µm). Representative of n=3 biological replicates. (**D,E**) qRT-PCR analysis showing the expression of p57 and miR221 in in clonogenic CSCs maintained under proliferative conditions, cultured in quiescence-inducing medium, or induced to re-enter the cell cycle by re-exposure to mitogenic stimuli. (D,E) Representative of n=3 biological replicates, each performed in technical triplicate. (**F**) Schematic representation of the in vitro differentiation protocol used to induce CSC differentiation into cardiomyocytes. (**G**) qRT-PCR analysis showing the expression of p57 and miR221 at Day 0 (D0), D4, D8, and D14 of the myogenic differentiation protocol. (**H**) Bar graphs showing the percentage of BrdU^pos^ cells in freshly isolated qCSCs+miR221 compared to qCSCs+miRNA control and in freshly isolated aCSCs+anti-miR221 compared to aCSCs+anti-miRNA control. (**I**) qRT-PCR analysis showing the expression of p57 in freshly isolated qCSCs+miR221 compared to qCSCs+miRNA control and in freshly isolated aCSCs+anti-miR221 compared to aCSCs+anti-miRNA control. (G-I) Representative of n=3 biological replicates, each performed in technical triplicate. (**J**) Bar graphs showing luciferase activity in qCSCs and aCSCs transfected with empty or p57_3’UTR miRNA reporter vector. Representative of n=3 biological replicates. All data are mean ± SD

To determine whether the reciprocal miR-221/p57 levels reflect CSC activity state per se --rather than an injury-specific signature--, we implemented an *in vitro* “quiescence–reactivation” assay analogous to previously used in other adult stem cells ^83^. Clonogenic CSCs were first maintained under proliferative conditions and driven into reversible quiescence by switching to quiescence-inducing medium and subsequently returned to cycling by re-exposure to mitogenic stimuli (Figure 7C).

Consistent with dynamic regulation by activity state, the marked decrease of BrdU incorporation in the quiescence-inducing medium (Figure 7C) was accompanied with p57 upregulation and miR-221 downregulation (Figure 7D,E). Conversely, upon reactivation, CSCs re-entered the cell cycle and p57 levels declined while miR-221 increased toward the proliferative level (Figure 7D,E). Thus, miR-221 and p57 track CSC cycling state in a reversible and opposite manner, consistent with a model in which miR-221 promotes CSC activation, at least in part, through repression of the quiescence driver p57.

To explore whether this axis also plays a role in CSC to CM differentiation pathway, we profiled miR-221 and p57 during *in vitro* cardiomyogenic differentiation of clonogenic CSCs (Figure 7F). Consistent with the activated state, clonogenic CSCs in proliferative conditions displayed high miR-221 and low p57 mRNA levels, and these levels were preserved during early aggregation into cardiospheres (Figure 7G), indicating that sphere formation per se does not revert the activation-associated miR-221^high^/p57^low^ profile (Figure 7G) suggesting that the CSC continue to multiply in the cardiospheres. In contrast, when CSC-derived cardiospheres were primed with myogenic morphogens to initiate CM differentiation ^57^, miR-221 transcripts progressively declined (Figure 7G), whereas p57 increased in a reciprocal manner (Figure 7G), consistent with release of miR-221–mediated repression and reinstatement of a p57-linked non-proliferative program. Notably, this reciprocal switch culminated with maximal p57 re-induction and minimal miR-221 levels coincident with the onset of spontaneous beating (∼day 10 through day 14) ^57^ in CM differentiation medium (Figure 7G), supporting a model in which the miR-221/p57 module not only tracks CSC activation/quiescence but is also dynamically modulated as the CSCs transition toward a cardiomyocyte fate.

Supporting this model, transient transfection of a miR-221 precursor into freshly isolated qCSCs efficiently induced their activation with a robust increase in BrdU incorporation (69.6±6.8% vs 8±1% in miRNA control) (Figure 7H) and a 30-fold downregulation of p57 mRNA expression (Figure 7I). The functional link between miR-221 and p57 repression was further confirmed by Luciferase assay (Figure 7J). In contrast, inhibition of miR-221 in aCSCs with a selective anti-miR reduced BrdU incorporation (13±4.6% vs 73±5.6% anti-miRNA control) (Figure 7H), accompanied by a ∼6-fold higher p57 expression as compared with anti-miRNA control (Figure 7I).

These findings demonstrate that the interplay of miR-221 and p57 regulates post-injury CSC activation through miR-221 induction which represses p57. Following repair, miRNA-221 is downregulated, p57 is upregulated and the CSCs return to the quiescent state.

### miR-221 enhances CSC regenerative capacity

To directly test the functional role of miR-221 in CSC regenerative competence, qCSCs^GFP^ and aCSCs^GFP^ were freshly isolated from control or DTA-injured Tg-Myh6^MCM^:R26^eGFPstop-DTA^ donor hearts, respectively, where the GFP tracks the fate of the transplanted cells and their progeny. After isolation, cells were subjected to a short recovery period under culture conditions optimized for viability and transfection efficiency while preventing the qCSCs from significantly reentering the cell cycle. qCSCs^GFP^ were then transfected with a miR-221 precursor, whereas aCSCs^GFP^ were transfected with an anti-miR-221, using matched miRNA controls. Because both reagents act transiently, with maximal efficacy within the first 72 h after transfection (*data not shown*), this approach (used here and also in the experiments described above in Figure 7H-J) allowed us to test whether temporary miR-221 upregulation or downregulation is sufficient to shift CSCs toward an activated or quiescent state, respectively, and whether such state transition is a prerequisite for full CSC regenerative competence.

*In vitro*, miR-221 overexpression in qCSCs^GFP^ markedly enhanced clonogenicity (9.3±1.5% vs 0.4±0.2% in miRNA control) and spherogenicity (7.8±1.9% vs 0.5±0.3%) (Figure 8 A-D) whereas miR-221 inhibition in aCSCs^GFP^ significantly reduced clonogenicity (1±0.5% vs 4±1% in anti-miRNA control) and spherogenicity (1.3±0.3% vs 4.5±1.3% in anti-miRNA control) (Figure 8 A-D). Thus, transient miR-221 gain of function drove qCSCs^GFP^ toward an activated phenotype, whereas transient miR-221 inhibition shifted aCSCs^GFP^ toward quiescence.

**Figure 8.**
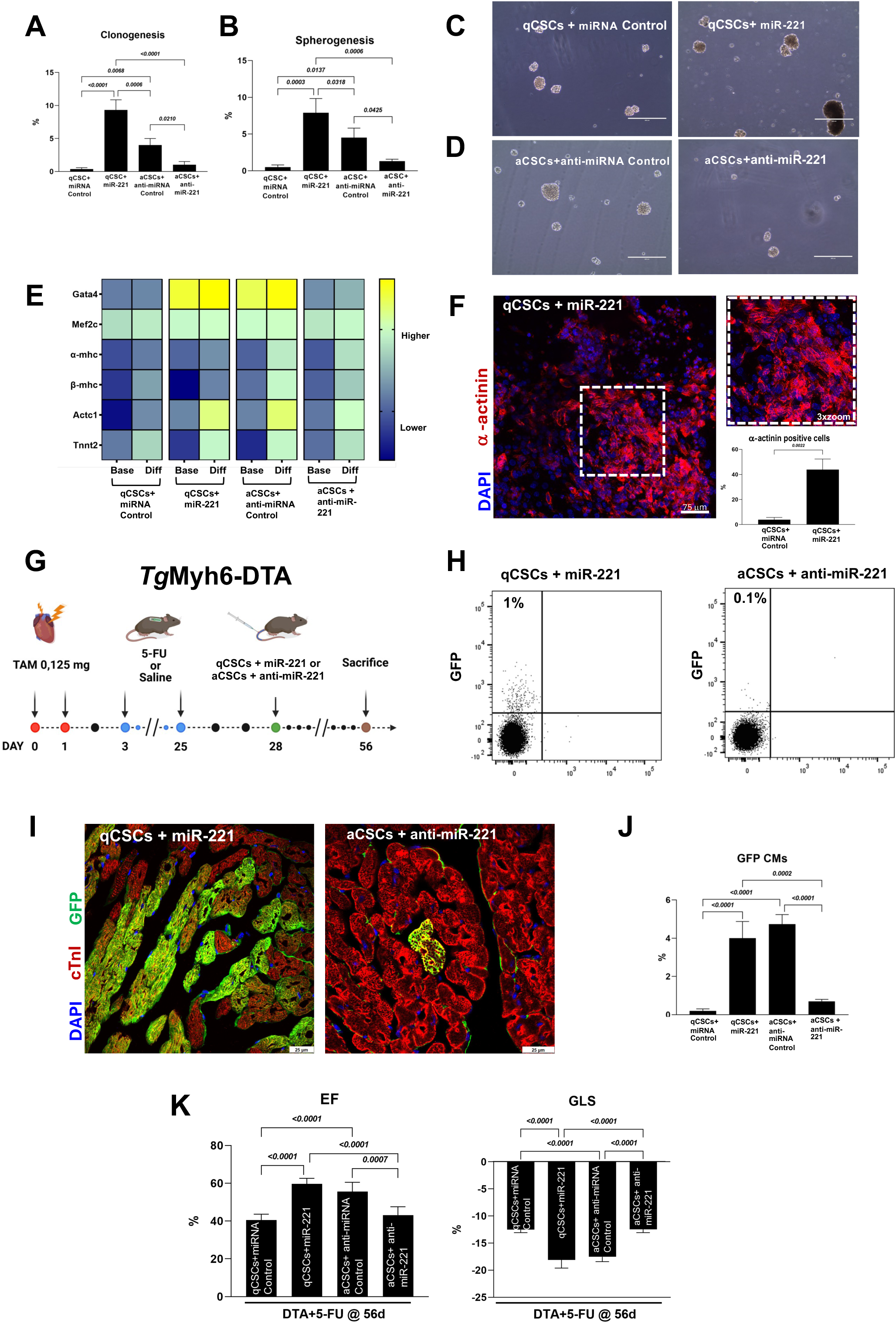
miR-221/p57 axis regulates cardiac stem cell proliferation and multipotency after injury. (**A,B**) Bar graphs showing the percentage of clones and cardiospheres in freshly isolated qCSCs transfected with a pre-miR miR221 (qCSCs^GFP^+miR221) and aCSCs^GFP^ transfected with anti-miR miR221 inhibitor (aCSCs^GFP^+anti-miR221) compared to their respective controls. Representative of n=3 biological replicates. (**C,D**) Representative light microscopy images of cardiospheres derived from qCSCs^GFP^+miR221 and aCSCs^GFP^+anti-miR221 compared to their respective controls. Scale bars = 400 µm. Representative of n=3 biological replicates. (**E**) Heatmap showing qRT-PCR analysis of the main myogenic genes in qCSCs^GFP^+miR221 and aCSCs^GFP^+anti-miR221 compared to their respective controls at baseline and after 14 days of myogenic differentiation in vitro. The color scale represents changes in Ct values (threshold cycle) relative to the GAPDH-normalized control. Representative of n=3 biological replicates, each performed in technical triplicate. (**F**) Representative confocal images and bar graph showing the percentage of α-actinin positive cells in qCSCs^GFP^+miR221 14 days after myogenic differentiation in vitro. (actinin, red; DAPI, blue nuclei, scale bars=75 µm). Representative of n=3 biological replicates. (**G**) Experimental design of the in vivo 5-FU-based approach used to ablate proliferating cardiac cells after TAM-induced injury in *Tg*Myh6-DTA mice with or without intravenous transplantation of qCSCs^GFP^+miR221 or aCSCs^GFP^+anti-miR221. (**H**) Representative dot plots showing the engraftment of qCSCs^GFP^+miR221 and aCSCs^GFP^+anti-miR221 into 5-FU-treated *Tg*Myh6-DTA recipient mice 28 days after TAM injection. Representative of n=3 biological replicates. (**I, J**) Representative confocal images and bar graph showing the percentage of GFP^pos^ CMs derived from qCSCs^GFP^+miR221 and aCSCs^GFP^+anti-miR221 after tail vein injection in DTA+5-FU-treated *Tg*Myh6-DTA recipient mice (GFP, green; cTnI, red; DAPI, blue nuclei, scale bars = 25 µm). Representative of n=3 biological replicates. (**K**) Bar graphs showing the percentage of the ejection fraction (EF) and the global longitudinal strain (GLS) in DTA+5-FU–treated *Tg*Myh6-DTA mice injected with qCSCs^GFP^+miR221 and aCSCs^GFP^+anti-miR221. The number of animals per group is reported in Supplementary Table 6. All data are mean ± SD

Notably, transient miR-221 overexpression also enhanced cardiomyogenic differentiation of qCSCs^GFP^ *in vitro*. Indeed, under pro-differentiation conditions, miR-221-treated qCSCs^GFP^ showed increased expression of key cardiac transcripts, including *Gata4, Mef2c,* and *Tnnt2*, together with a marked increase in α-actinin-positive cells displaying organized sarcomeric structures compared with miRNA control (43.8±8.4% vs 3.8±1.8%) (Figure 8 E,F). Conversely, miR-221 inhibition markedly impaired cardiomyogenic differentiation of aCSCs^GFP^, as shown by reduced expression of the same cardiac genes and by a lower proportion of α-actinin-positive cells (8±3% vs 47±5.6% in anti-miRNA control) (Figure 8 F). Together, these findings indicate that miR-221 functions as a molecular switch, rather than as a continuously acting driver, in regulating CSC state. Induction of miR-221 in qCSCs is sufficient to trigger exit from quiescence and acquisition of an activated, cardiomyogenically competent phenotype, whereas transient miR-221 inhibition in aCSCs is sufficient to restore a quiescence-like state. Once triggered, these state transitions appear to become self-sustaining. Accordingly, miR-221 overexpression in qCSCs likely initiates activation, at least in part through repression of p57, allowing the newly activated cells to remain competent to respond to cardiomyogenic cues and progress toward differentiation. Conversely, miR-221 inhibition in freshly activated aCSCs resets the cells toward a quiescence-like state, thereby reducing their ability to undergo efficient cardiomyogenic differentiation. Overall, our data indicate that CSC activation is not merely associated with proliferation but is also a prerequisite for full cardiomyogenic and regenerative competence.

To determine whether these state changes also operate *in vivo*, miR-221-modified qCSCs^GFP^ and aCSCs^GFP^ were injected intravenously into *Tg*Myh6-DTA recipient mice with DTA+5-FU-induced cardiomyopathy (Figure 8 G). Twenty-eight days after transplantation, miR-221-overexpressing qCSCs^GFP^ showed significantly greater cardiac engraftment than qCSCs^GFP^ miRNA control (1±0.3% vs 0.1±0.1%), whereas anti-miR-221-treated aCSCs^GFP^ showed reduced engraftment compared with aCSCs^GFP^ anti-miRNA control (0.1±0.1% vs 0.9±0.2%) (Figure 8 H). Importantly, miR-221-overexpressing qCSCs^GFP^ generated substantially more GFP-positive CMs than qCSCs^GFP^ miRNA control (4±0.9% vs 0.2±0.1% of total CMs), while anti-miR-221-treated aCSCs^GFP^ showed a marked reduction in GFP-positive CM formation compared with their anti-miRNA control counterparts (0.7±0.1% vs 4.7±0.5%) (Figure 8 I, J). Functionally, mice receiving miR-221-overexpressing qCSCs^GFP^ exhibited cardiac recovery comparable to that observed after transplantation of aCSCs^GFP^ miRNA control, whereas anti-miR-221-treated aCSCs^GFP^ and qCSCs^GFP^ miRNA control failed to support functional improvement (Figure 8 K, Supplementary Figure 6 and Supplementary Table 6).

Altogether, these findings identify miR-221 as a central regulator of CSC activation and regenerative competence. By repressing p57, miR-221 promotes exit from quiescence and acquisition of an activated state that is required for robust clonogenic, cardiomyogenic, and regenerative potential. Conversely, inhibition of miR-221 in activated CSCs drives them back toward a quiescence-like state and markedly reduces their myogenic and regenerative capacity. Thus, miR-221-dependent CSC activation emerges as a prerequisite for full CSC regenerative competence and effective CSC-mediated cardiac repair.

## Discussion

Several major conclusions emerge from this study. First, the adult murine heart is not an entirely cell-static organ but retains substantial regenerative capacity when CM loss is diffuse and the coronary circulation and supporting myocardial architecture remain intact. Second, functionally defined adult CSCs reside within the CD45-negative/CD31-negative population expressing c-kit and/or Sca-1. Although their clonogenicity, self-renewal, and multilineage differentiation potential have been established in previous studies, the present work directly addresses their most contested property by demonstrating robust cardiomyogenic competence *in vitro* and, most importantly, *in vivo*. Third, activation of this resident CSC compartment supports replacement of up to approximately one tenth of the left ventricular CMs within one month. Fourth, the limited or absent contribution of CSCs reported in previous studies reflect, at least in part, the use of injury models that disrupt the vascular, stromal, and extracellular-matrix environment, such as myocardial infarction, as well as intrinsic limitations of the employed genetic lineage-tracing strategies. Fifth, although the present study does not formally exclude cardiomyocyte renewal through proliferation of pre-existing differentiated CMs, any such contribution was below the level detectable under our experimental conditions. Finally, the CSC response to cardiomyocyte loss follows a reversible quiescence–activation–differentiation–return-to-quiescence program regulated by the reciprocal miR-221/p57 axis: p57 maintains CSC quiescence, whereas injury-induced miR-221 upregulation suppresses p57 and promotes cell-cycle entry and regenerative competence; subsequent miR-221 downregulation and p57 re-expression accompany cardiomyogenic differentiation and restoration of CSC quiescence after repair.

Cardiomyocyte regeneration and renewal have long been a matter of intense debate ^5, 6^. Although multiple studies have reported new CM formation in the adult heart in response to injury and to cumulative wear and tear with aging, this process is regarded as negligible in adult cardiac biology, physiology, and disease ^5–7, 35^. Although the post-natal mouse heart has been shown able to regenerate its apex ^84, 85^, CMs become terminally differentiated and unable to re-enter the cell cycle soon after birth ^86, 87^. The prevailing cell-static view of the adult heart is further supported by the fact that the mouse heart is reported to have a CM renewal rate of ∼1% per year ^61, 88, 89^, and a similar rate has been estimated by radiocarbon dating for the human heart ^1, 2^ despite their very difference size and lifespan. All these data implied that, from cradle to grave, life depends on the cohort of CMs present shortly after birth, but which are inexorably decreasing by the wear and tear of the uninterrupted heartbeat.

The most striking finding of this study is the magnitude and rapidity of the endogenous regenerative response. Following the acute loss of approximately 15% of left ventricular CMs, the adult heart generated new CMs equivalent to nearly one tenth of the LV CM compartment within only 28 days, with restoration of myocardial architecture and complete recovery of ventricular function (Figure 3). These findings further suggest that the limited repair observed after myocardial infarction may not simply reflect an intrinsic inability of the adult heart to generate new CMs, but also the simultaneous destruction of the vascular, stromal, and niche components required to sustain an effective regenerative response. Thus, the adult myocardium appears capable of quantitatively meaningful regeneration, but the expression of this capacity is critically dependent on the nature of the injury and on preservation of the tissue environment in which regeneration must occur.

Of note, Hume et al. ^90^ recently identified a marked increase in mitotic and cytokinetic activity among small mononuclear cardiomyocytes in the peri-infarct border zone of the adult human heart after myocardial infarction, with locally enriched populations reaching approximately 20%. These findings support the existence of an intrinsic, injury-responsive cardiomyocyte regenerative program in humans and are consistent with our results.

### The Cardiac Stem Cells (CSCs)

A small sub-population of all c-kit+ adult myocardial cells (<10%) has been repeatedly shown to be self-renewing, clonogenic and multipotent *in vivo* and *in vitro* ^12, 23, 25, 55^. When properly stimulated these cells give origin to the four main myocardial cell types: spontaneously beating cardiomyocytes, smooth and endothelial vascular cells, and fibroblasts ^12, 23, 25, 55^. Demonstration that the progeny of one of these cells expanded *in vitro* (a clone) when transplanted, either intramyocardially or through the systemic circulation, homes and nests into damaged myocardium where it generates new CMs and micro vessels, justifies calling them cardiac stem cells (CSCs) ^12, 23, 25, 55^. Their presence has been identified in all mammalian species studied, from mice to human, by a plethora of independent investigators ^12–15, 17–24, 91^. Nonetheless, <1% of all myocardial c-kit+ cells are CSCs; the rest are blood-borne cells and endothelial cells and their precursors ^23, 55^. Until now, no CSC-specific membrane marker has been found to allow for their identification and direct isolation. Therefore, their isolation depends on cumbersome exclusion of endothelial and blood lineage cells from the c-kit+ cell population ^23, 55^. Among the CD45^neg^/CD31^neg^/c-kit^pos^/Sca-1^pos^ up ∼20% show myogenic potential *in vitro,* which raises significantly *in vivo,* as shown here.

### The CSCs home, nest, multiply and differentiate in the damaged myocardium

When delivered through the coronary circulation, CSCs, nest within, and differentiate inside damaged myocardium. After administration through either the systemic ^25, 55^ or coronary circulation ^92^ in models of myocardial injury, transplanted CSCs selectively home and nest within the injured myocardium, reconstitute the depleted CSC compartment, and generate new cardiomyocytes and micro-vessels with consequent structural and functional recovery. A key component of this process is the activation of the SDF-1/CXCR4 chemotactic axis: myocardial injury induces local SDF-1 expression, while CSCs express CXCR4, enabling directional migration toward the damaged tissue, trans-endothelial recruitment, and retention within the permissive regenerative niche ^25^. We had further shown that interference with this pathway markedly impairs CSC homing and myocardial integration, identifying SDF-1/CXCR4 signaling as a central mechanism of CSC trafficking to sites of injury ^25^. In parallel, the HGF/c-Met and IGF-1 signaling pathways provide complementary motogenic and survival cues, with HGF promoting CSC migration and tissue invasion and IGF-1 supporting survival, expansion, and regenerative competence after engraftment ^92^. Consistent with this biology, intracoronary delivery of IGF-1 and HGF in large-animal infarction models activates endogenous cardiac stem/progenitor cells, increases their accumulation in infarct and border zones, enhances the formation of new myocardium and microvasculature, reduces infarct size, and improves ventricular performance ^92^. Thus, the available evidence indicates that successful CSC transplantation through the bloodstream depends on a coordinated sequence of chemokine-guided homing, adhesion and extravasation, niche-dependent nesting followed by cardiomyogenic and vasculogenic differentiation, rather than on simple passive retention. The injured myocardium provides the instructive molecular gradients required for this process to occur at therapeutically meaningful levels.

### The Controversies

As it often happens in response to paradigmatic shifts, the initial broad interest elicited by the first description of the CSCs was rapidly followed by questions and skepticism about their nature and physiological significance. Firstly, doubts about their existence were based on the lack of any specific marker for their isolation from the bulk of myocardial c-kit+ cells. Tests of different mixed populations of myocardial c-kit^pos^/Sca-1^pos^ cells failed to show a CM fate ^93–95^. Curiously, some groups described cardiomyocyte precursor cells using their preferred marker (c-kit, Sca-1, MDR-1, Abcg2, Isl1 and so forth), with the puzzling consequence that the adult heart, for decades considered as lacking stem cells or myogenic progenitors, ended up with an astonishing number of apparently different CSCs ^12–15, 17–24^. Subsequent work has shown that the different markers were either co-expressed by the same cell or at different physiological, differentiation or activation stages of the same cell lineage ^96^. Yet, the lack of a CSC-specific identifying membrane marker remains a source of confusion. Second, the physiological import of the CSCs was challenged because, clearly, they are unable to significantly regenerate the massive segmental myocyte losses caused by an AMI ^5–7^. This objection ignored the fact that endogenous tissue regeneration, even in the most regenerative tissues, repairs wear and tear damage or satisfies the physiological cell turnover of the related tissue but not the tissue losses produced by occlusion of segmental arteries ^8^. Thirdly, the *coup-de-grace* to the nascent field of CSC biology and myocardial regeneration came from attempts to genetically determine the cell fate of the CSCs and to identify their progeny^97–99^. Using either c-kit or other non-specific genes expressed in the CSCs, Cre-lox (and other similar site-specific recombinase systems) cell-fate mapping studies failed to show a significant contribution of the c-kit^pos^ cells to new CM formation during development, in adulthood or after injury, with concomitant denial of the existence of any resident adult cardiomyocyte progenitor cell ^30, 31, 36–40, 100^. However, because the generation of new CMs in adulthood could no longer be denied, the failed cell-fate mapping studies were swiftly followed by reports claiming that the main/sole source of adult neo-cardiomyogenesis was the replication of adult and terminally differentiated myocytes! ^61, 101, 102^.

Genetic cell-fate mapping approaches, despite their very significant contribution to stem cell biology and tissue/organ morphogenesis and regeneration, have significant limitations depending on their setup ^103–105^. To generate reliable data three conditions must be satisfied: i) the recombinase should be inserted into a gene that identifies the cell whose fate is to be mapped without affecting its expression or producing a null allele with subsequent haploinsufficiency of the targeted gene; ii) the recombinase should efficiently recombine and label all/most target cells avoiding underestimation of its differentiation potential; iii) the recombination system should not be ectopically expressed leading to an overestimation of the progeny of the cell tracked. Unfortunately, none of the genetic cell-fate mapping reports claiming a minimal/non-existing contribution of CSCs to CMs so far published meets the above criteria ^106^. As a key example, the first report that c-kit expressing cardiac cells do not or only minimally contribute cardiomyocytes ^39^, ignored the fact that the CSCs represent only <1% of all c-kit+ myocardial cells and its setup (Cre inserted into the first intron of c-kit) produces a c-kit haploinsufficiency which suppresses the cardiomyogenic potential of CSCs and, in addition, fails to recombine the majority of them ^55^.

A related limitation applies to dual orthogonal site-specific recombination strategies designed to distinguish pre-existing cardiomyocytes from all nonmyocytes, which have been reported as the ultimate proof to negate existence and role of CSCs ^105^.

In contrast, the present experiments test CSC cardiomyogenic cellular and functional potential: intravascular transfer of the progeny of a single CSC into DTA-injured, 5-FU-treated decompensated hearts reconstituted the host CSC compartment and generated new cardiomyocytes, leading to structural and functional recovery (Figure 5). These findings recapitulate, confirm and extend our previous work demonstrating that CSCs are necessary and sufficient for myocardial regeneration following isoproterenol-induced injury ^25^.

### Regulation of CSC quiescence and activation, a pre-requisite for CSC full regenerative competence

In multiple adult organs, regenerative competence relies on the capacity of resident adult stem cells to exit quiescence, transiently expand, and reconstitute the damaged parenchyma in a spatially and temporally restricted manner ^8–11^. Hematopoietic stem cells, skeletal muscle satellite cells, intestinal crypt stem cells, and epithelial stem cells ^8–11, 70, 73, 107^ among others, follow this pattern: quiescence preserves the pool, injury- or stress-induced cues unlock a controlled proliferative and differentiation program, and its return to quiescence prevents exhaustion of the pool and prevents neoplasia ^70, 72–74, 108^. The data shown here that diffuse CM loss triggers a quantitatively meaningful regenerative response, place the adult heart within the broader framework of injury-responsive, stem-cell–dependent tissue repair. The adult myocardium is no longer exceptional but behaves as a regulated regenerative tissue rather than a purely post-mitotic organ.

The identification of the p57/miR-221 axis as a molecular enforcer of the CSCs quiescence and determinant of their activation provides both mechanistic insight and a direct target for potential regenerative strategies. p57^Kip2^ is a canonical cyclin-dependent kinase inhibitor that enforces quiescence and restricts cell-cycle entry across diverse stem cell compartments^78, 79^. Its downregulation is required for controlled activation of tissue-resident progenitors, whereas its persistent expression maintains the quiescent, protected state ^78, 79^. Also, p57^Kip2^ is a known critical regulator of embryonic myocardial cell differentiation and, together with pRB, is a barrier to adult cardiomyocyte proliferation ^109–111^. Here we show that injury-induced miR-221 upregulation in CSCs is coupled to p57 suppression, resulting in cell-cycle entry and CSC expansion. The reverse transition is required for CSCs to enter an intermediate state permissive for acquisition of a CM-committed phenotype, in which miR-221 is downregulated and p57 re-upregulated. This is consistent with prior evidence implicating miR-221 in the promotion of proliferation and survival in other cell types, including vascular and progenitor populations, through repression of cell-cycle inhibitors ^112, 113^. In CSCs, however, this quiescence-suppression transition acts not as an oncogenic switch, but as a tightly regulated module that converts a quiescent regenerative reserve into an active reparative/regenerative pool in response to myocardial injury, before being reset to promote CSC return to quiescence while its progeny progresses into terminally differentiated CMs.

## Author contributions

Conceptualization: E.C., B.N-G., F.M., and D.T.; methodology and investigation: E.C., M.S., A-J.S., C.S., L.P., C.Q., N.S., A.DiC., G.C., F.M.; funding acquisition: E.C., D.T.; supervision: A.DeA., G.M.E-H, K.U.; writing –original draft: E.C., F.M., B.N-G., and D. T.; writing – review & editing: E.C., B.N-G., and D. T.

## Data availability

Further information and requests for materials, resources and reagents as well as any additional information required to re-analyse the data reported in this paper should be directed to and will be fulfilled by the lead contact, Dr. Daniele Torella.

